# Scan-o-matic: high-resolution microbial phenomics at a massive scale

**DOI:** 10.1101/031443

**Authors:** Martin Zackrisson, Johan Hallin, Lars-Göran Ottosson, Peter Dahl, Esteban Fernandez-Parada, Erik Ländström, Luciano Fernandez-Ricaud, Petra Kaferle, Andreas Skyman, Stig Omholt, Uros Petrovic, Jonas Warringer, Anders Blomberg

## Abstract

The capacity to map traits over large cohorts of individuals – phenomics – lags far behind the explosive development in genomics. For microbes the estimation of growth is the key phenotype. We introduce an automated microbial phenomics framework that delivers accurate and highly resolved growth phenotypes at an unprecedented scale. Advancements were achieved through introduction of transmissive scanning hardware and software technology, frequent acquisition of precise colony population size measurements, extraction of population growth rates from growth curves and removal of spatial bias by reference-surface normalization. Our prototype arrangement automatically records and analyses 100,000 experiments in parallel. We demonstrate the power of the approach by extending and nuancing the known salt defence biology in baker’s yeast. The introduced framework will have a transformative impact by providing high-quality microbial phenomics data for extensive cohorts of individuals and generating well-populated and standardized phenomics databases.

## INTRODUCTION

While our ability to detect genetic variation has improved tremendously (Koboldt et al. 2013; Mardis 2013), our capacity to rapidly and accurately map the phenotypic effects of this variation in large population cohorts – phenomics – has not made comparable gains (Houle et al. 2010). This is true also for microbes, where estimation of fitness components defines the key entrance point when searching for the concerted effects of causative genetic variation. As the majority of genetic or environmental effects on net fitness are modest in size (Thatcher et al. 1998; Wagner 2000), this demands access to measurement methodology capable of capturing subtle differences.

Micro-colony analysis estimates within-population variations (Levy et al. 2012) but is low throughput for differences between populations. Miniaturization of liquid cultures allows automation and highly accurate measurement of population density (Warringer and Blomberg 2003, 2014) but its associated costs in time and manpower are too high to allow massive scale-up. Tagging of strains with DNA sequence barcodes and monitoring tag frequencies in complex strain mixtures allows high throughput but at lower accuracy and at high initial investments (Giaever et al. 2002; Winzeler et al. 1999). Better cost-efficiency can be achieved using ordered arrays of microbes that are cultivated on solid media as colonies whose area sizes are estimated using cameras (Costanzo et al. 2010; Kvitek et al. 2008; Bean et al. 2014; Lawless et al. 2010). The set-up is attractive in that microbes are surveyed in a growth mode that better resembles their natural state, but the environment can rarely be maintained constant across plates. This leads to large spatial biases and false results. Spatial bias correction is a formidable challenge (Baryshnikova et al. 2010) because colonies exchange nutrients and toxins with the local environment and affect each other. Accentuating problems, 3-dimensional colony population size is typically approximated by measuring the often poorly correlated 2-dimensional colony area (Pipe and Grimson 2008). Furthermore, growth tends to be estimated from a single or few measures (Tong et al. 2001; Tong et al. 2004). As any particular colony size can be reached via an endless number of very different growth paths (Fig 1A), this unavoidably creates false positives and negatives as well as wrong conclusions.

**Figure 1.**
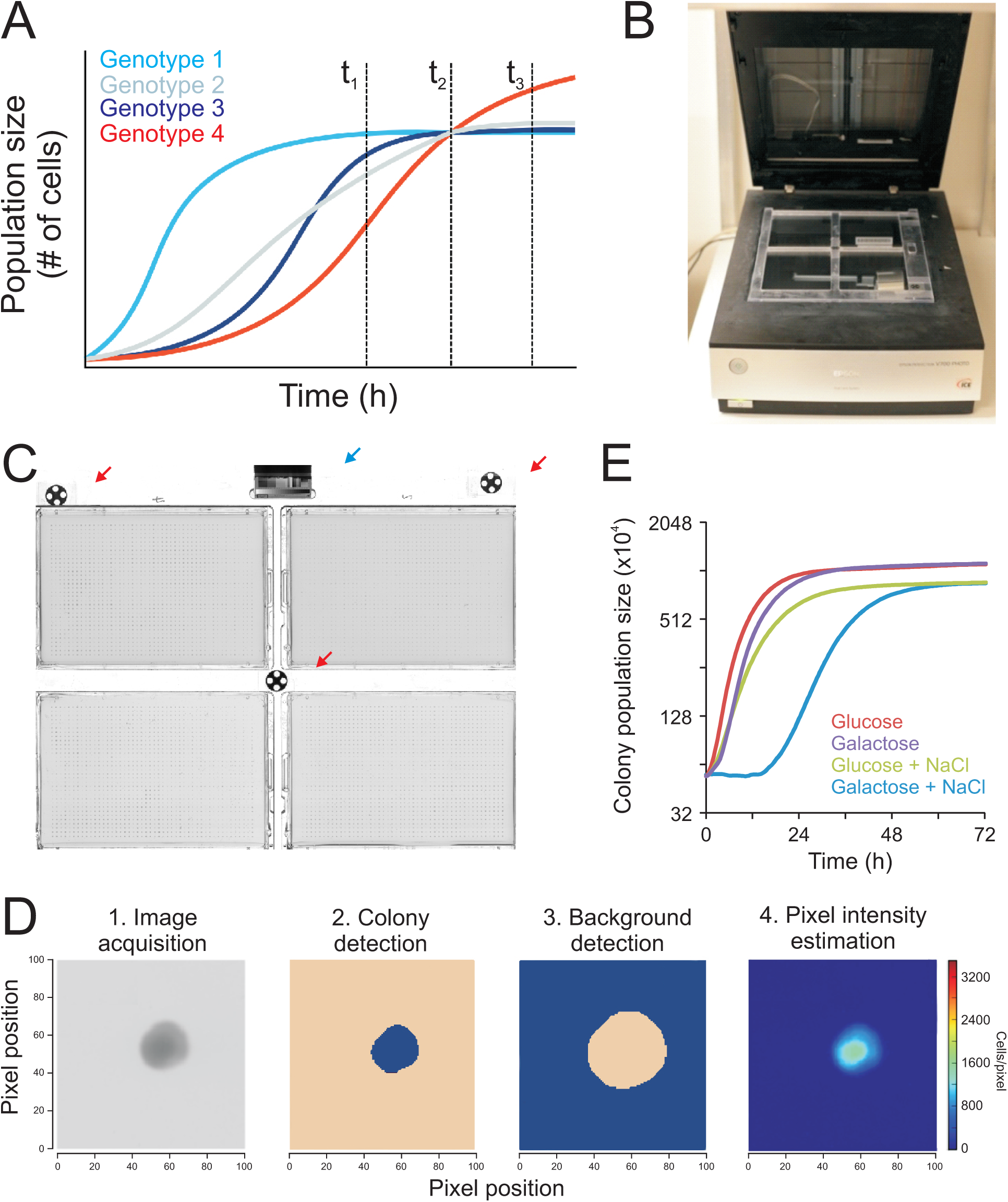
– A novel framework for high-resolution microbial phenomics. A) Single time point measures at different stages of growth (vertical broken lines) provide wildly diverging views of the relative growth performance of strains, alternatively scoring all genotypes as identical (t_2_), genotype 1 as superior (t_1_) and genotype 4 as superior (t_3_). This illustrates how effects on distinct aspects of the cellular physiology of different genotypes can be falsely called in the absence of dynamic growth data. B) Scan-o-matic is physically based on high quality desktop scanners and fixation of four solid media plates on top of the scanning surface using custom-made acrylic glass fixtures. C) Four solid media plates – each containing 96 to 1536 microbial colonies – are fixed on each scanner by the glass fixture. Three black & white orientation markers (red arrows) on the fixture allow precise tracking of pixel positions and a fixed transmissive grey scale calibration strip (blue arrow) permits between-scans and scanners calibration of pixel intensities. D) Colony population size is extracted from raw images in a multi-step procedure, proceeding from raw image, to probable colony (blob; blue) detection and segmentation, local background definition (blue) with a safety margin to colony, and estimation of cells as pixel intensity as compared to background. Colour intensity = 0 (dark blue) – 1500 (turquoise) cells per pixel. E) Colony population growth curves obtained by cultivating genetically identical WT colonies in four environmental contexts and measuring colony population size in 20 minute intervals. Y-axis is on log(2) scale.

The near-term goal of microbial phenomics is therefore to achieve the accuracy of liquid microcultivation in a solid media cultivation mode. To reach this goal the 3-dimensional topology of microbial colonies must be captured at high spatial and temporal resolution and reduced into accurate estimates of colony population sizes while exhaustively compensating for local environmental variations. Here we present a novel microbial phenomics methodology dubbed Scan-o-matic that overcomes the above hurdles and demonstrate its utility on the eukaryotic model organism baker’s yeast, *Saccharomyces cerevisiae.*

## RESULTS

### A novel framework for high-resolution microbial phenomics on solid media

To develop a high-resolution microbial growth phenomics framework for the average microbiology lab, we established a pipeline based on high-quality mass-produced desktop scanning technology (Fig 1B). The scanners allow transmissive light capture, enabling precise estimates of cell counts in microbial colonies (see below). Scans are initiated at pre-programmed intervals by means of a power manager that switches off scanner lamps immediately after scanning. This drastically reduces spatial bias from exposure of colonies closer to the lamp parking position to excessive light and temperature. Four solid media microcultivation plates, each containing from 96 to 1536 colonies, fit in each scanner (Fig 1C). Given initial instructions, Scan-o-matic completes the entire data acquisition to growth feature extraction pipeline autonomously (Fig S1, S2). Plates are fixed within an acrylic glass fixture where their precise positioning is automatically detected using orientation markers (Fig S3). Each fixture contains a transmissive grey-scale calibration strip to compensate for variations in pixel intensities between scans that otherwise represent a major obstacle for robust estimation of colony population size (Fig S4). Scan-o-matic segments images and robustly localizes colonies, avoiding misclassification from dust particles, plastic deformations or scratches (Fig 1D, S5, S6). Colony intensities are compared to local background pixel intensities and converted via a calibration function into cell counts per pixel (Fig S7). These are summed over the colony to produce precise estimates of colony population size. High frequency of estimates translates into well-resolved growth curves that are quality controlled with a semi-automated user interface (Fig S8).

Cultivation over a range of environmental contexts shows the vast majority (>99.5%) of growth curves to have the classical sigmoid shape, to be minimally affected by noise and systematic artefacts, and to have distinct environment specific properties (Fig 1E). Following a lag phase of variable length, early growth was found to be near exponential, and in unstressed environments it corresponded to a raw population size doubling time of 2.0h (Coefficient of Variation, CV = 4.9%, *n* = 1536). Carbon limited populations growing on simple sugar sources reached distinct stationary phases when the carbon was depleted. Curve shapes were independent of pinning format (384 or 1536 colonies) (Fig S9A). Final cell number increased at lower colony pinning density presumably due to reduced competition for carbon and energy. However, initial population size was smaller in the denser formats as the smaller pin-heads deposited fewer cells. The net effect was that the 1536 format provided a longer growth span than the 384 format (mean of 5.1 vs. 4.9 doublings, p<10^−8^), faster growth and more stable estimates of the growth rate (Fig S9B).

The trade-off between image resolution and image acquisition time currently makes high-quality data beyond the 1536 format unattainable. Each scanner handles four plates and each computer controls three scanners; thus, even our prototype arrangement with five computers and 15 scanners is capable of running more than 92,000 individual growth experiments in parallel, using the 1536 format. This throughput is orders of magnitude better than what can be achieved by liquid microcultivation.

### Enhanced measurement precision and accuracy of colony population size and growth rate

Highly time-resolved growth data offers decisive conceptual advantages since distinctive physiological states can be analysed (Fig 1A). To test whether our approach also offers technical advancements, we compared its precision and accuracy to that of the current microbial phenomics standard on solid media: measures of 2-dimensional area covered by the colony at a single point in time. When considering 1536 pinning format plates containing genetically identical cultures (wildtype, WT), colony population size growth curves contained less random noise than their colony area counter parts (Fig 2A). Using the standard error of the regression at the time of maximal growth as a measure of random curve noise, the population size growth curves were roughly two-fold as robust as the colony area growth curves (Fig 2B). This rather drastic reduction in random noise was expected, because the 2-dimensional colony area measures per definition contain less and have less resolved information. This difference in information content increases as growth proceeds to reach a maximum at stationary phase entry. Here, colony growth, measured as image area covered by the colony, corresponded to a single doubling on average, while colony population size typically corresponded to more than four doublings (Fig 2A). Thus, the critical measure of precision, random noise as a fraction of signal strength (CV), vastly favours colony population size growth curves.

**Figure 2.**
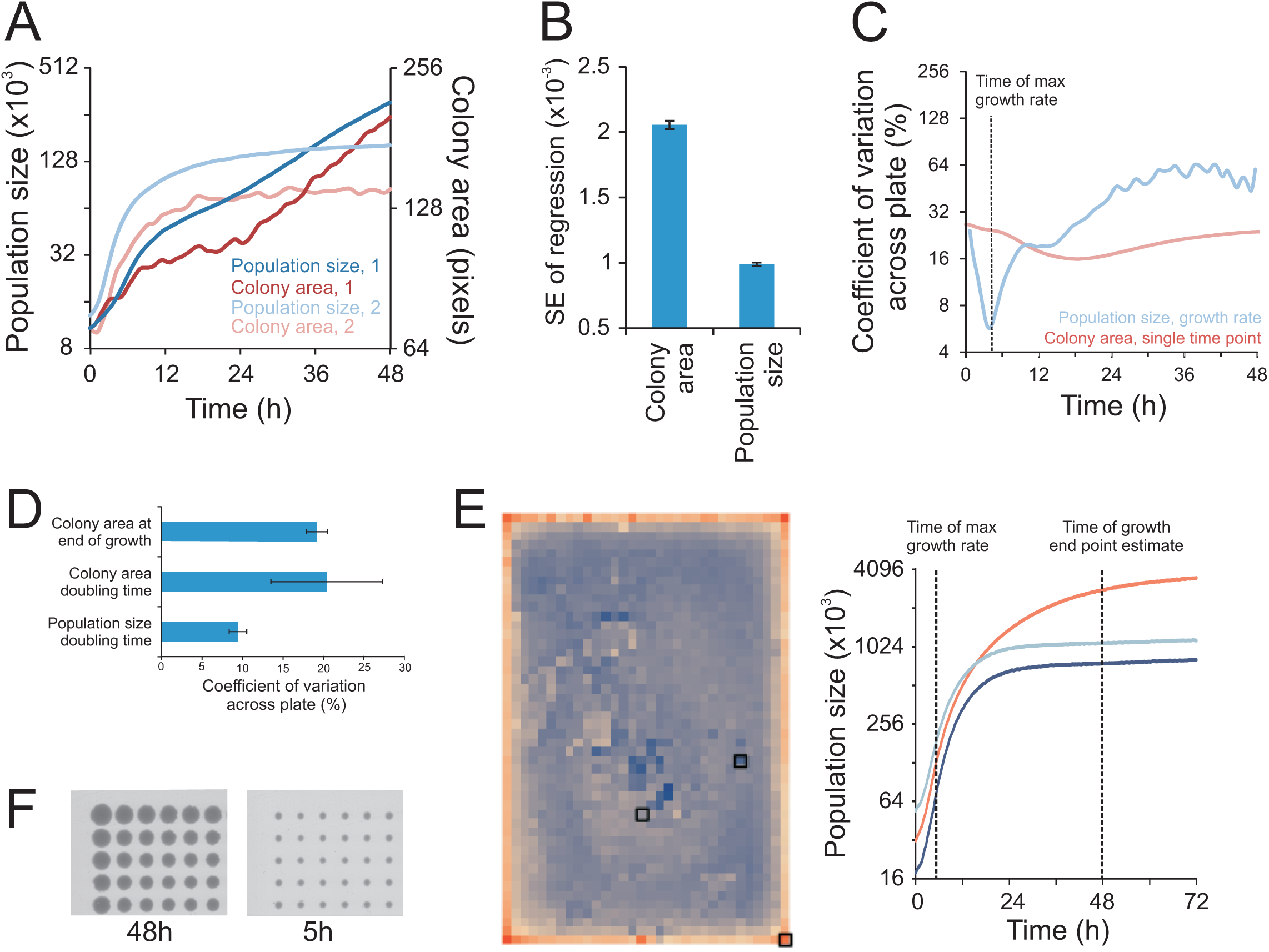
– Dramatically enhanced measurement accuracy in microbial phenomics. A-B) Comparing random noise in growth curves based on either colony area or colony population size. A) Growth curves of two sample colonies, labelled 1 and 2. Y-axes are on log(2) scale. B) Estimating growth curve noise in the critical section of the curve when growth is maximal. Noise was measured as the standard error of the regression corresponding to the highest slope. Mean of 1536 genetically identical WT growth curves in an unstressed environment is shown. Error bars = SEM. C-D) Accuracy (sum of random noise and systematic bias) over a plate, measured as the coefficient of variation across 1536 genetically identical WT colonies. C) Accuracy as a function of time, for colony area size and colony population size growth rate. A single plate of unstressed populations is depicted. Y-axis is on log(2) scale. D) Accuracy for single measure of colony area at end of growth (48h), colony area growth rate and colony population size growth rate. The mean of four plates with different stresses (2% glucose and 2% galactose, with and without 0.85M NaCl) is shown. Error bars = SEM. E) *Left panel:* Visual representation of the edge effect for colony area at end of growth (48h). Each square corresponds to one of 1536 genetically identical (WT) colonies grown in absence of stress. Colour intensity shows colony area, dark blue = 500 and red = 1800 pixels. Bold squares indicate colonies highlighted in the right panel. *Right panel:* Population size growth curves for colonies indicated in the left panel. Curve colour matches the colour of the highlighted squares in the left panel. Time points of maximal growth rate and for 48h measures are indicated (broken lines). F) Raw image of a corner section of a 1536 plate with genetically identical (WT) colonies growing in absence of stress, at 48h and at the time of maximal growth rate (5h).

Precision provides an incomplete picture of technical achievement by overlooking systematic bias. Consideration of the total measurement error for a plate as the variance over genetically identical colonies divided by their mean showed accuracy to shift dramatically depending on growth state (Fig 2C). Accuracy was initially low for both area and population size, primarily due to large variation in the robotic delivery of cells, but also due to larger influence of randomness when pixels representing each colony were few (initially ~100). The total measurement error for colony area on a plate steadily decreased to reach a minimum late in the growth phase. Thereafter it rapidly increased as competition intensified, favouring growth of outer frame colonies with fewer neighbours. Variation in growth rates obtained from colony population size began similarly high, but dropped dramatically to reach a very low minimum well below single point measures in early exponential growth phase, when growth rates are maximal. We therefore extracted maximal growth rates as population size doubling time estimates as the most stable, key feature of growth curves.

Considering six diverse environments, population size doubling times were estimated with about twice the accuracy of single measures of colony area at the end of growth (Fig 2D). This increase in accuracy is partly due to the lack of competition between colonies at the time at which growth is maximal and thus lower spatial bias (Fig 2E, F). The avoidance of colony competition effects represents not only a quantitative but also a qualitative advancement by averting confounding comparisons of strains in different physiological states (Fig 1A).

### Comprehensive removal of spatial bias by reference-surface normalisation

Assuming the common, but naïve, null hypothesis approach that all systematic bias has been removed by the experimental design, we would expect ≈ 5% false positives for each of the above single genotype (WT) tests (*α* = 0.05, Student’s t-tests). However, even after extensive technical optimization and using the most robust measure of growth (population size doubling times) to minimize error, we scored many more (~10x) false positives than chance expectation (Fig 3A). This suggested substantial systematic bias to remain. The change in error-to-signal ratio over time (Fig 2C) indicated two distinct types of errors to dominate population size doubling time estimates: an initial bias from the number of cells deposited on plates that decreases in magnitude as colony population size increases, and a later, increasing bias that derives from increasing competition between neighbouring colonies (Fig 2E). Of these, the former is the most important, as reflected in a strong correlation between population size doubling times and initial population size (Fig 3B). However, bias from initial population size only explained about half of the error, as shown by the remaining false positives after normalizing the population size doubling times to the initially deposited number of cells (Fig 3A).

**Figure 3.**
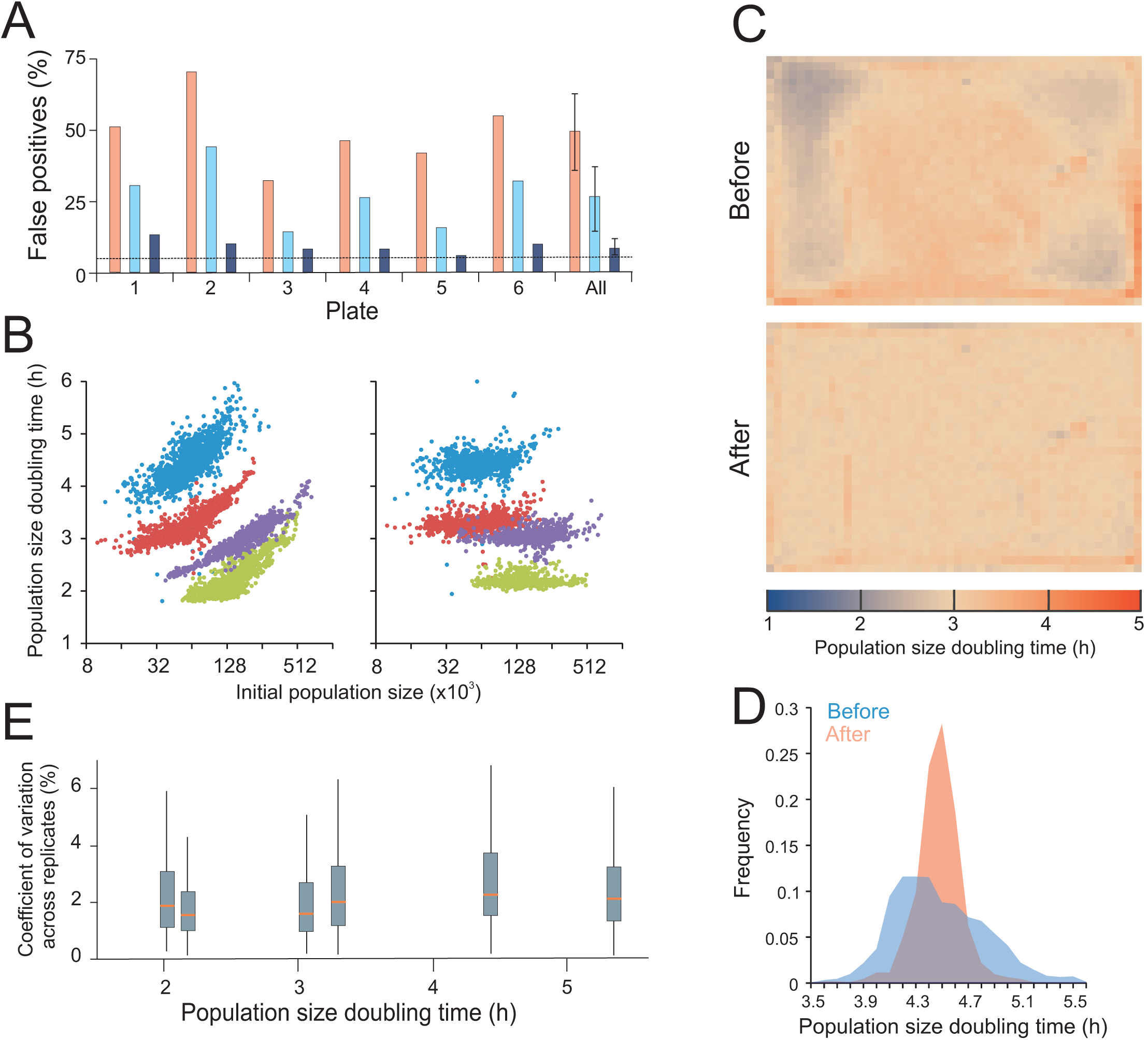
– Comprehensive removal of spatial bias in microbial phenomics. A) Fraction of false positives due to spatial bias within plates with genetically identical colonies (WT). Each plate corresponds to one distinct environmental challenge. On each plate, population size doubling times of immediately adjacent colonies (excluding every 4^th^ position that was used a control position) were statistically compared to those of non-adjacent colonies using a one-sample Students t-test (H_0_ = zero difference, α=0.05). Assuming all variation to be random, i.e. no spatial bias, the random expectation is 5% false positives (broken line) at this significance cut-off. Any excess of false positives corresponds to spatial bias. Pink bars = before normalization, light blue bars = after normalization to initial population size, dark blue bars = after reference grid normalization. “All” indicates the mean of false positives over all six plates with error bars = SEM. B) Population size doubling time as a function of initial population size. Left panel = before normalization, right panel = after reference grid normalization. All individual estimates over four of the six genetically homogeneous 1536 plates with different environmental challenges (2% glucose and 2% galactose, with and without 0.85M NaCl) are shown. C) Spatial bias is removed by reference grid normalization. Genetically identical reference colonies are pinned into every fourth colony position (lower right position in every tetrad of positions), creating a matrix of 384 control colonies on which a normalization surface of population doubling times is based. The local normalization surface is subtracted from each observation. Upper panel = distribution of population size doubling times of 1536 genetically identical colonies across a plate, before normalization. Each square corresponds to a colony position. Colour indicates population size doubling time. Lower panel: As upper panel, but colour represents population size doubling times after reference grid normalization. Normalization was achieved by designating each 4^th^ position a control position. See also Fig S10. D) Frequency distribution of population size doubling times, before and after reference grid normalization, in a sample plate (blue experiments in B, 2% galactose + 0.85M NaCl. E) Box plot showing variation in population doubling time estimates (y-axis, CV between adjacent colonies) after normalization within each of the six genetically identical but environmentally distinct experiments in A, as a function of plate mean population doubling times (x-axis). Red line = median CV for all groups of adjacent colonies on plate, box = inter quartile range (mid 50%) of CVs, whiskers = complete range of CVs.

Topology and strength of the spatial bias varied dramatically between plates and environments to produce complex patterns (Fig 3C, S10 - *before* panels). The major tendency in the bias was a positive correlation between neighbours that directly contrasted with the competition-for-resources model believed to explain bias at later stages. To meet this challenge and to stringently normalize for complex and unpredictable spatial bias of unknown origin, we replaced every fourth position on the plates with isogenic controls, creating an array of 384 reference positions (Fig S11). This defines an evenly spaced 2-dimensional array of control colony growth rates that by interpolation reflects local spatial variations over plates with high fidelity. Subtraction of the spatial control-surface of growth rates from actual estimates provided position-normalized measures. This normalization removed the correlation of population doubling time to initial population size (Fig 3B), and it considerably reduced variation across plates (Fig 3C, S10). Critically, it also resulted in a false positive rate that approached random expectations by being an order of magnitude lower than the one obtained without spatial normalization (Fig 3A). Spatially normalized population doubling times were approximately normally distributed, a fundamental requirement for application of standard parametric statistics (Fig 3D). Spatial normalization also completely removed the correlations between relative error (CV) and signal strength (population doubling times) that otherwise would complicate statistical treatment (Fig 3E). After spatial normalization the total measurement error in an unstressed environment was around 2% of the signal, approaching the 1.5% achieved with state-of-the-art microcultivation in liquid media (Warringer et al. 2003; Warringer et al. 2011). Thus, although some spatial bias that is not fully captured by neighbouring controls does remain, this reference-surface normalization protocol permits a measurement accuracy that almost matches the best that can be achieved with lower throughput approaches.

### Scan-o-matic extends and nuances the salt biology of baker’s yeast

Gene-by-salt interactions have previously been called by scoring NaCl-sensitive gene deletion strains with state-of-the art microcultivation, and the underlying biology extensively detailed (Warringer et al. 2003). To evaluate the recapitulation of established knowledge, we compared our new method with previous microcultivation experimental calls of salt-sensitive mutants in the complete yeast gene deletion collection.

Normalising to growth defects in absence of any stress (Fig 4A), we identified a large number of salt-sensitive gene deletions. Many of these, such as those encoding Hog1 and Pbs1 that activates the osmo-response (Hohmann 2002) or the transcription factor Crz1, corresponded to proteins well known to control salt tolerance and exhibited growth curves on solid and liquid media that were very similar (Fig 4A, B). Among the salt-sensitive mutants identified in this genome-wide screen, 14 biological processes were enriched. Many of these, e.g. ion homeostasis, ion transport, response to osmotic stress, and endocytosis and vacuolar transport (Fig 4C), have previously been shown to be of importance during salt exposure (Warringer et al. 2003).

**Figure 4.**
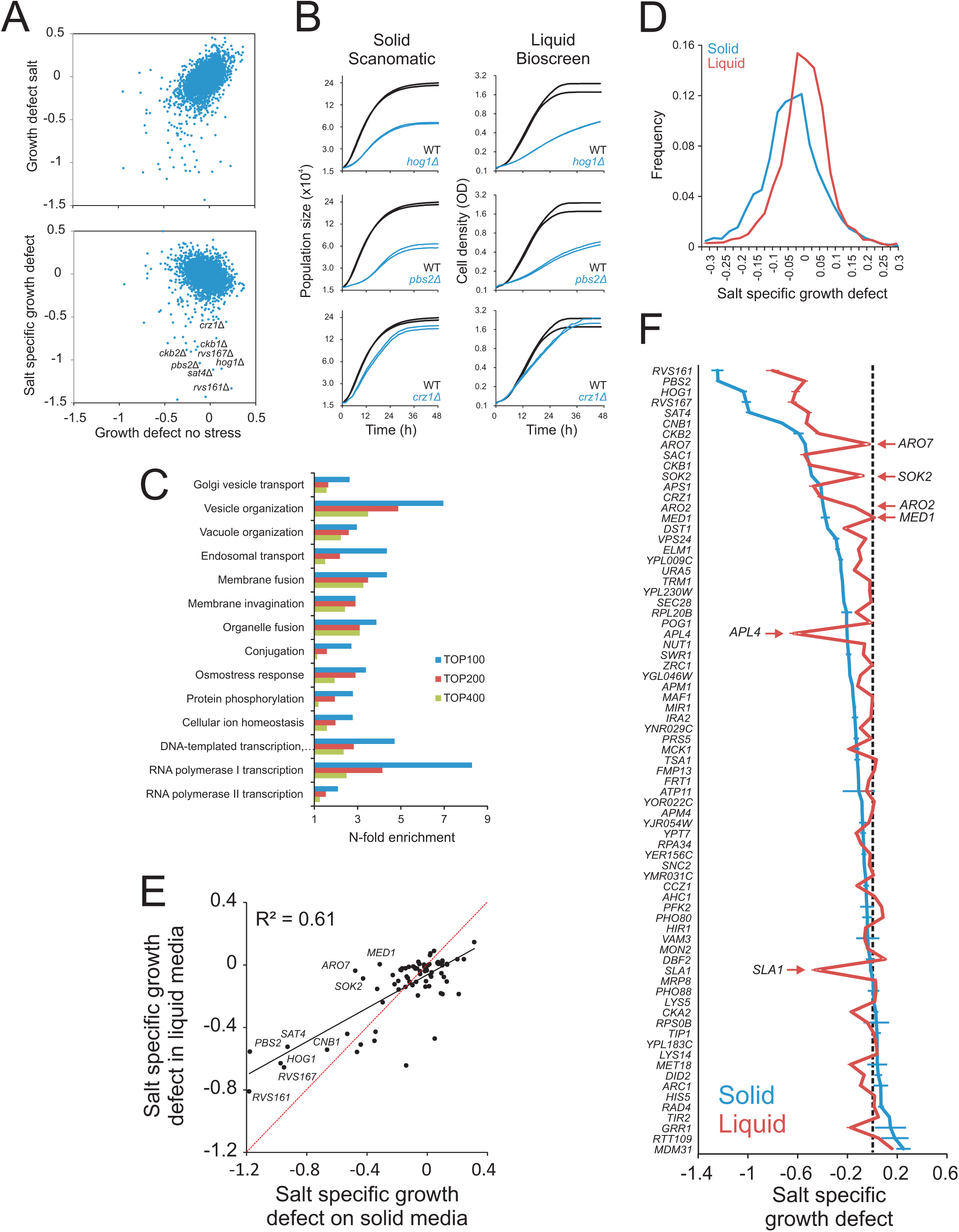
– Recapitulating and extending the known salt biology with Scan-o-matic. The haploid *MAT***a** yeast deletion collection was cultivated in 2% glucose, with and without 0.85M NaCl. Log(2) population doubling times relative the control surface of WT controls were extracted. Negative values represent growth defects. A) *Upper panel:* Comparing relative population size doubling times of the yeast deletion collection in presence and absence of NaCl. *Lower panel:* Colony population doubling times in NaCl were normalized to corresponding measures in absence of NaCl, estimating NaCl-specific growth effects. These were plotted as a function of relative population doubling times in absence of NaCl. Little correlation remains. B) Growth dynamics of three sample deletion strains (n=2) with NaCl specific growth defects, in Scan-o-matic and during liquid microcultivation. C) Functions enriched (Fisher’s exact test, FDR *q*<0.05) among top 100, 200 and 400 most salt sensitive deletion strains in Scan-o-matic. Cut-offs approximately correspond to relative growth defects larger than −0.27, −020 and −0.15. D) Frequency distributions of salt-specific deletion strain growth effects, obtained by solid substrate cultivation in Scan-o-matic and by liquid microcultivation in a Bioscreen C (Warringer and Blomberg 2003) E-F) A subset of 70 deletion strains were re-cultivated in absence and presence of 0.85M NaCl at high replication, using Scan-o-matic (solid; n=24) and liquid (n=6) microcultivation, respectively. Re-cultivations were performed in parallel, removing all conceivable systematic variation beside cultivation method. E) Salt-specific growth defects in Scan-o-matic and liquid microcultivation regimes. Regression (black, Pearson *R*^2^ is indicated) and 1:1 lines (red) are shown. F) Deletion strains were ranked based on salt-specific growth effects during solid substrate cultivation and salt specific growth effects were plotted. Error bars = SEM.

However, the distribution of gene-by-salt interactions was substantially wider using our new set of protocols, with a distinct shoulder towards salt sensitivity (Fig 4D). Thus, at any given signal strength and at any given significance stringency, more interactions were identified on agar compared to liquid microcultivation. The amplification of signal strengths of traits was evident also with regard to growth in absence of stress, suggesting the increase of phenotypic variation on solid media to be a general phenomenon (Fig S12A). Notably, we also found that genes that were required for salt tolerance during liquid microcultivation were in many cases irrelevant for salt tolerance when scored on agar (data not shown). This suggests that differences in salt-specific genotype-by-environment interactions were due to different demands on cells dispersed in a solution compared to cells fixed in a colony structure.

To exclude batch and temporal bias and stringently test the influence of cultivation method on the gene-by-salt interactions, we re-screened the same subset of genotypes with both cultivation methods using identical media, random positioning and parallel screening. We found correlation between cultivation methods to be only intermediate (*r*^2^ = 0.5 - 0.6), both in the presence and absence of salt (Fig 4E, S12B, S12C). Roughly 10% of the phenotypic variation could be explained by technical noise, meaning that the remaining 30–40% derived from phenotypic effects imposed by cultivation method differences.

Both cultivation methods called the most important salt-defence regulators, like signalling components Hog1 and Pbs1, Rvs161 and Rvs167 that reorganizes the action cytoskeleton during Na^+^ stress (Balguerie et al. 2002; Lombardi and Riezman 2001), the regulatory subunits Ckb1 and Ckb2 of the casein kinase that regulates Na^+^ extrusion (Glover 1998), the regulatory subunit of calcineurin Cnb1 plus its downstream transcription factor Crz1 that controls Na^+^ efflux at current pH (Stathopoulos and Cyert 1997), and the cation extrusion regulator Sat4 (Mulet et al. 1999) (Fig 4E). However, most mutants were salt sensitive in only a single cultivation regime. Thus, despite an abundance of amino acids in the media, the absence of either Aro2 or Aro7, required for synthesis of aromatic amino acids, was central to salt tolerance only during colony agar-growth, as were the removal of the RNA pol II component Med1 or the inhibitor of pseudohyphae formation, Sok2 (Fig 4F). In contrast, presence of the clathrin coated vesicle component Apl4 and the actin cytoskeleton associated Sla1 were critical to salt tolerance only for cells reproducing dispersed in a solution. Overall, 81% of re-tested gene deletions differed significantly (Student’s t-test, FDR *q*<1%) in salt tolerance between solid media and microcultivation growth, the majority (68%) of phenotypes being amplified on solid media. Thus, whereas our new approach recapitulated the essence of established salt biology wisdom, it also highlighted the large influence of cultivation approach on both the quantitative and qualitative aspects of phenotypes.

## DISCUSSION

Scan-o-matic as currently implemented drastically improves and sets a new standard in microbial growth phenomics but does not exhaust opportunities for future improvements.

Further reductions in random noise would require changing from 8- to16-bit image depth, increased scanning frequency or enhanced image resolution, all of which are associated with trade-offs. 16-bit image depth increases file size, challenging the logistics of data storage and analyses, and is poorly supported. Scanning frequency and image resolution are in a direct quality trade-off as enhanced image resolution reduces scanning speed and thereby increases exposure to radiation, heat and dehydration from light. This is a serious concern because exposure to intense light severely impedes growth (Logg et al. 2009), however, the light-sensitive hog1 and pbs2 mutants where not affected by the scanning frequency currently employed (data not shown).

Reductions in bias require meticulous attention to experimental design or to posterior correction procedures. Early spatial bias derives primarily from systematic variations in numbers or physiological states of cells deposited, the latter emerging due to transfer of metabolically distinct central or peripheral members of pre-culture colonies. To ensure that the normalization surface compensates for such bias, controls and experiments should originate from the same pre-culture plate. Late spatial bias derives from variations in nutrient access and toxin exposure that depend on media thickness and homogeneity as well as number and size of metabolically active colonies in the local environment. Standardization of plate casting procedures are thus essential as are media buffering because cell secretion of acidic metabolites otherwise drives patch-wise pH variations (Fig S13) affecting local growth. Earlier published posterior correction procedures have used spatial variations in the performance of experimental colonies themselves and assumed that spatial bias mainly manifests as an edge effect that can be compensated for in a row/column wise manner (Baryshnikova et al. 2010). Adoption of this approach as an additional level of spatial bias normalization is unlikely to improve accuracy, first because use of experimental data both as correction input and as final output creates unsound statistical dependencies and artefacts and second because the edge effect is not a major driver of bias at the time of maximal growth rates. Due to its dramatic nature, with plateaus of low and high bias respectively and their separation by sharp fault lines, comprehensive posterior removal of the remaining spatial bias is extremely challenging.

Condensation of population growth curves into estimates of maximal growth rates makes poor use of the vast trove of accumulated data. Time before net growth commences (growth lag) and the total gain in population size (growth efficiency), are additional fitness components (Cooper 1991) capable of driving adaptation (Ibstedt et al. 2015). Higher noise in the beginning and end of growth means that more attention to the experimental design and analysis procedure is needed before robustness can be achieved also for these fitness components. Beyond lag and efficiency, colonies rarely maintain maximal growth rates over extended periods of time. In addition, growth often becomes multiphasic (Warringer et al. 2008). A single growth rate estimate, maximal growth rate, will thus not fully capture the true dynamics of growth. Future more exhaustive use of growth information will help ensure that the introduced Scan-o-matic framework provides truly high-quality microbial phenomics data for extensive cohorts of individuals to generate well-populated, highly resolved and standardized databases.

## MATERIALS AND METHODS

### Physical arrangement of Scan-o-matic

Scan-o-matic is based on high quality mass-produced desktop scanners (Fig 1A) that are controlled by power-managers (Fig S1A). Images are acquired using SANE (Scanner Access Now Easy) (Mosberger 1998) using transmissive scanning at 600 dpi, 8-bit grey-scale and a scan area extension that captures four plates per image (Fig 1B). Plates are fixed in custom-made acrylic glass fixtures with orientation markers ensuring software recognition of fixture position (Fig 1B). Fixtures are calibrated to scanner by a fixture calibration model (Fig S2B, S4). Pixel intensities are standardized across instruments using transmissive grey scale calibration targets (Fig S4). Scanners are maintained in a 30°C, high humidity environment and kept covered.

### Scan-o-matic software, image acquisition and analysis

Scan-o-matic is written in Python 2.7 and can be installed from https://github.com/local-minimum/scanomatic/wiki. Matplotlib is used for graph production. Numpy, Scipy and Scikits-Image is used for computation and analysis (van der Walt et al. 2011; van der Walt et al. 2014). Experiments are initiated from a web-interface (Fig S2A), 7 minutes being the minimum time interval between scans (20 min as default) and 96, 384 or 1536 pinned plates as allowed formats. Each series of scans is analysed in a two-pass process. The first-pass is performed during image acquisition (Fig S1B). Using a fixture calibration model (Fig S2B) and orientation markers (Fig S3A), the plate and transmissive grey scale calibration strip positions are identified (Fig S3B). The transmissive grey scale calibration strip area is trimmed (Fig S4A) and segment pixel intensities are compared to manufacturer’s supplied values (Fig S4B). In the second-pass analysis (Fig S1C), a virtual grid is first established across each plate so that grid intersections match the centres of likely colonies (Fig S5). At grid intersections, the local area is segmented, colonies defined relative the local background and pixel intensities of both are determined, compared and converted into actual cell number estimates (Fig S6, S7). Colony image areas were determined as the number of pixels included in each colony definition.

### Extracting, evaluating and normalizing population growth rates from smoothed growth curves

Raw growth curves are smoothed, first using a median filter that removes local spikes and then using Gaussian filter that reduces remaining noise. Initial colony population size, maximal population size doubling time, time of extraction of the maximal population size doubling time, error of the linear regression that underlies extraction of the maximal doubling time, and the growth curve fit to an initial value extended version of the classical Chapman-Richard model are extracted from smoothed growth curves. The latter three are used to flag poor quality growth curves that are visually inspected for potential rejection (~0.3% were here rejected).

Every 4^th^ experimental position was here reserved for internal isogenic controls. Doubling times of controls were used to establish a reference surface as follows. Controls with extreme values were removed. Remaining control positions were then used to interpolate a normalization surface that was smoothed, first with a median filter to exclude any remaining noisy measurements, and then with a Gaussian smoothing to soften the landscape contours. For each colony, the log(2) difference between the observed doubling time and the doubling time of the normalization surface in that position was calculated. When multiple plates were included in an experimental series (Fig 4), we first normalized for between-plates bias by shifting the mean of the non-spatially normalized controls on each plate to match the mean over all plates. To call gene-by-salt interactions (Fig 4) we normalized growth rates in NaCl by subtracting growth rates in a no stress environment.

### Wet-lab experimental procedure

Solid media plates were cast with 50mL of Synthetic Complete (SC) agar medium buffered to pH 5.8 with 2% glucose or 2% galactose as carbon source and with and without 0.85M NaCl. Two strain layouts were used: 1) all colonies being diploid BY4743 reference strain (Brachmann et al. 1998) (Fig 1–3) 2) colonies corresponding to single yeast gene knockouts of the haploid BY4741 deletion collection (Giaever et al. 2002), with WT control colonies interleaved in every fourth position and n=3 replicates of each strain in juxtaposition (Fig S11). For the confirmation experiment (Fig 4E-F, S12B-C) the same procedure was employed, but at high (n=24 replicates). The reference liquid media experiments (Fig 4E-F, S12B-C) were as previously described (Warringer et al. 2011).

## SUPPLEMENTARY INFORMATION

Supplementary Materials and Methods

Supplementary figure S1-S13

Supplementary movie S1

## ACKNOWLEDGEMENTS

We thank Olle Nerman and Mats Kvarnström for much appreciated statistical and analytical support and Charlie Boone for access to strains. Financial support from the Swedish Research Council (325–2014–6547 and 621–2014–4605) and from the Carl Trygger Foundation (CTS 12:521) to JW, from the Swedish Research Council FORMAS to AB, from the Slovenian Research Agency (P1–207 and L2–1112) to UP and PK and from and LABEX SIGNALIFE (ANR-11-LABX-0028–01) to JH is acknowledged.

## SUPPLEMENTARY MATERIALS AND METHODS

### 1. Physical arrangement, Hardware and Hardware Control of Scan-o-matic

#### 1.1 Pinning and plates

Scan-o-matic is based on the use of a pinning robot, which allows for swift and reliable transferring of colonies from source plates to target plates. Here, the RoToR robot (Singer LTD, UK) was used. The plastic, disposable pinning pads used by the robot are fixed but not fully rigid 2D matrices of pins that only allow transfers in multiples of 96. Supported plate formats are 96, 384, and 1536. Scaling up between colony formats is done by interleaving repeated pinnings according to default, as well as custom-made programs (see below). The latter is specifically designed to minimize measurement noise and bias using the Scan-o-matic normalization principles. It is critical to create an even agar surfaces, both for target and source plates. Even surfaces decreases initial spatial variation in population size, due to uneven deposition of cells, and decreases local variations in access to nutrients later in the growth curve. Even surfaces are achieved by having an absolutely even and level surface when casting plates. To avoid batch bias between plates, it is imperative to pour exactly 50mL of medium into each plate, to pour into the same position of each plate and to maintain temperature at pouring constant over time.

#### 1.2 Scanner

Scan-o-matic utilizes mass-produced technology available to an average microbiology lab at a reasonable cost. Three Epson Perfection V700 PHOTO scanners (Epson corporation, UK) are connected via USB to each controlling computer (Fig. S2A). The power supply of each scanner is controlled individually and independently from a computer using a GEMBIRD EnerGenie PowerManager LAN (Gembird Ltd, the Netherlands) (Fig S2A). Each experiment is run as a separate autonomous process, but scanner power supply toggling is coordinated. This ensures that malfunction in one scanner does not affect other scanners connected to the same computer. Experiment processes do not require the user interface to continue running for their operation. Images are acquired using SANE (Scanner Access Now Easy), a standardized Linux interface for scanning (Mosberger 1998). Scanning is performed using transmissive scanning with the transparency unit (TPU) fixed at 600 dpi, 8 bit grey scale. The V700 has two TPU modes. The officially documented TPU mode is unable to support scanning of the full scanner area, and can only capture two plates in each image. To enable simultaneous capture of four plates and allow sufficient throughput, we updated the back-end for Epson scanning in SANE to support a second TPU mode. This mode captures the full scanning area – sufficiently large to contain four plate images. The second TPU mode is available from Ubuntu 13.04 onwards. For that reason, we recommend using the long term release Ubuntu 14.04. The scanner firmware lacks control options for shutting down the scanning lamp rapidly after image acquisition, which causes it to stay on for > 15 minutes after scanning. Extensive light exposure leads to drying out of the solid agar medium, contraction of the agar and colony repositioning. It also causes excessive and spatially biased variation in radiation, water evaporation and temperature. Effects combine to create severe spatial bias and variations in scanner sensor properties over time, the latter leading to peak shifts in image histograms and loss of measurement precision. Due to EU regulations regarding stand-by mode in electronic appliances, scanners have been physically modified by insertion of a plastic wedge behind (and thus short-circuiting) the power button. Scanners are therefore ready to scan when given power. To allow instant shutting down of the scanner lamp after image acquisition, scanner power supplies were connected to a GEMBIRD EnerGenie Power Manager LAN, a programmable power manager with an internal web-server. The Scan-o-matic software automatically logs into the power manager over the local area network, turning on and off sockets via its web-interface, as scanners are needed. Further complicating scanner control, individual Epson V700 scanners lack unique identifiers. Thus, there is no default way of distinguishing multiple scanners connected to the same computer. Scan-o-matic instead tracks the provision of power to scanners, ensures that scanners are provided with power in a serial manner (i.e. they are fired up serially but scanning can and does proceed simultaneously), and links power provision to assignment of USB ports.

#### 1.3 Computers

Physically, Scan-o-matic is run on standard desktop computers with the following suggested minimum requirements: Linux operating system, 2 GB RAM, Intel Core i3, 2TB storage. The analysis segment of Scan-o-matic, i.e. all section excluding visualisation and quality control of growth curves, also works on Mac.

##### 1.4 Plate Fixture

Each scanner has an individual, custom-made CNC milling machine cut acrylic glass fixture with four slots, each slot designed to take one plate. Each slot has pegs along its edge to ensure that plates are inserted with a fixed orientation, an otherwise common human mistake. Fixtures have three orientation markers (Fig S4A). These are used to triangulate the position of the plates and the calibration target within the fixture, both of which are initially unknown to the computer. The three orientation markers allow precise identification of plate and calibration target positions, thus avoiding any bias arising from slight movement of the fixture between scans. Each fixture also has a slit where a scanner calibration target, containing a transmissive scale, is inserted (Fig S5A). The calibration target is necessary to normalize pixel intensities between scanners, as well as to account for fluctuations in sensor sensitivity and lamp temperatures between scans in a scanner in an experiment series. The latter issue is greatly reduced with the above described power manager, but remains a potential source of bias. In the present work, the transmissive scale section of Kodak Professional Q-60 Color Input Target (Kodak company, USA) was used as a calibration target. However, the framework also supports calibration targets from SilverFast (LaserSoft Imaging AG, Germany).

### 2. Scan-o-matic software

Scan-o-matic is written in Python (2.7). The user interface is currently being migrated from GTK 2.0 to a more flexible and maintainable HTML5/JavaScript front-end Flask server solution. The modules NumPy, SciPy and Scikits-Image (van der Walt et al. 2011; van der Walt et al. 2014) are used for computation and analysis (http://www.scipy.org/). Information about full suite of dependencies required is bundled with the application and is available at: https://github.com/local-minimum/scanomatic/wiki/Installation.

### 3. Scan-o-matic analysis pipeline

#### 3.1. Design of a custom-made robotic pinning program for the Scan-o-matic pipe-line

A primary objective in the Scan-o-matic design is incorporation of a reference grid of control positions to produce a dense and evenly distributed matrix of controls. This matrix allows establishment of a three-dimensional growth normalization function that is used for reduction of spatial bias by adjusting the measurements of each colony to local variations in the function. Such variations occur in every single experiment due to a complex combination of factors that includes: i) variation of initial amount of cells deposited by the pins ii) uneven distribution of nutrients and stressors in the medium iii) medium thickness iv) spatial variations in light exposure, temperature, radiation and water evaporation v) interactions with neighbouring colonies by competition for shared resources and secretion of toxic compounds. Furthermore, if controls are kept constant over time, they also allow for normalization of drift in plate, scanner, and medium properties over time. However, Scan-o-matic does not explicitly require the use of this particular experimental design and imposes no particular constraints on the arrangement of strains. In the standard experimental design with a dense carpet of controls, two source plates are used, denoted A (or “strain source plate”) and B (or “control source plate”) (Fig S12). Strain source plate A contains the strains for which the experiment is designed. The control source plate B is a uniform plate with identical colonies of the control strain in all positions. The strain source plate A is pinned to three different, but neighbouring, offsets onto the target plate, C. The control source plate B is only pinned once onto a single position of the target plate, placing a control colony in the empty fourth position in each tetrad of positions. Thus, the target plate, C, will contain three replicates of each strain of interest, with the fourth position in each tetrad of positions containing a control strain replicate. Observe that a merging of the two source plates A and B into a target plate C should occur in the pre-culture, or before the pre-culture step, such that C is not equal to the actual experiment plate placed in the scanner. Otherwise, control position spatial bias will not equal experiment position bias and the normalization process loses much of its power.

#### 3.1 Fixture calibration

In each scanner, four experimental plates are precisely positioned using the custom-made plastic fixtures (Fig 1A, B). Each scanner should have its own fixture, avoiding potential noise and bias arising from fixtures with slightly different properties being shifted between scanners. The fixture fixes each plate so that spatial positioning is constant and precise during an experiment. Once per each fixture-scanner combination, a spatial calibration model that describes the layout of the fixture is created using the Scan-o-matic fixture calibration user interface (Fig. S3B). On a scanned image of the fixture, the number of fixture orientation markers, plate positions, and the position of the transmissive scale calibration strip are marked by interactive click-and-drag mouse maneuvers. The type of calibration target is selectable from a drop down menu. Scan-o-matic uses this fixture calibration as a model to automatically section images of said fixture into their meaningful components, discarding irrelevant image information and thereby greatly reducing the complexity of the image analysis, computation time, and errors.

#### 3.2. Initiating a Scan-o-matic experiment

Experiments are initiated from the Scan-o-matic user interface in which scanning parameters are set and meta-data identifying the experiment is supplied (Fig S3A). Parameters that are set are: scanner, fixture, experiment duration (from 14 minutes to 7 days, but typically 2–3 days), time interval between consecutive scans (from 7 to 180 minutes but typically 20 minutes), and the pinning density of each plate (96, 384, 1536, 6144). The user is also asked to document the contents of the experiment. The experiment is initiated from the interface and will run autonomously until completion. Results are automatically stored and analysed as described below. Each experiment can be interrupted if needed. This will end image acquisition and start analysis of the truncated project. A power outage will result in a terminated experiment, but the user can resume action when power returns as a new project and later join the two projects into a complete one.

#### 3.3 Image Acquisition

Image Acquisition is described under hardware above (see 1.2).

#### 3.4 Image Analysis

Image analysis is run as a two-pass procedure in order to optimize computational resources. The first pass is run while the experiment is progressing. The second pass is launched after the experiment is completed and is run in reverse, from the last time-point images to the first.

##### 3.4.1 First Pass Analysis

First pass analysis, referred to as “project compilation” in the user interface, is responsible for setting up the information needed for growth estimations. Fig S2B shows an overview of the first pass analysis. To enable positioning and annotation of image features in a manner that is consistent over images, orientation markers are used (Fig 1B).

###### 3.4.1.1 Localizing orientation markers in each image

The first step in the first pass analysis localizes orientation markers within each individual image. In essence, each image, *I*, is thresholded with a heuristically set pixel value of 127 (value range midpoint) resulting in a black and white image. A set, *C*, that stores centre coordinates of orientation markers is created. The thresholded image is convolved using the fast Fourier transformation function with a black and white representation of the fixture orientation marker, *M* (Fig S4A). This evaluates, for each pixel of the image, how well the local context of that pixel matches a representation of a fixture orientation marker. Iteratively, with the number of iterations corresponding to the number of fixture orientation markers in the fixture model, the strongest signal position is selected and chosen as the centre position of that fixture orientation marker, *c*. This centre position is stored appended to *C* and later used to calculate the origo for image features localization vectors (Fig S4B). A strong signal is obtained when the majority of the adjacent pixels correspond to the expected marker two colour pattern. To avoid reporting of the same marker twice, the local area around c, corresponding to the size of the convolution kernel, is set to one less than the minimum of the convolution surface. The process of marker localization is repeated on the modified surface until all markers have been found.

###### 3.4.1.2 Matching image orientation marker positions to fixture calibration model orientation marker positions

Second, fixture orientation marker coordinates of each image are compared to the corresponding coordinates of the fixed calibration model. The positions of image features are adjusted so that they perfectly agree with the model. In essence, all permutations, *P*, of marker coordinates, *C*, of the analysed image are evaluated for their distance, *d*, to the coordinates in the calibration model, *R.* The permutation with the smallest sum-value is assumed to describe the correct marker pairing. The offset vector between the two-dimensional centre of masses of the two coordinate systems are calculated and applied to all the plate and calibration scale coordinates of the model such that their coordinates are adjusted to relevant coordinates for the current image. This allows exact determination of the positions of plates within each image, independent of if the fixture used were damaged or moved during the experiment. It also greatly reduces the complexity of localizing grid of colonies that the pinning robot dispensed, allowing for gridding to be done once for all images. As a consequence, the pixels used to estimate the local background of each colony will be consistent throughout the experiment. This removes a source of noise. Since colonies will be largest towards the end of the experiment, it also makes gridding easier and more reliable. It should be noted that the robustness of both the gridding and the grey scale detection (see below), due to these solutions, is high enough that the plate areas and the grey scale area, i.e. the position of the fixed calibration target, of the fixed calibration model only needs to be a rough estimate of its actual areas.

###### 3.4.1.3 Image sectioning and pixel intensity calibration

For each image, the four plate containing areas and the calibration target containing area are sectioned out (Fig S4B). The image section defined as containing the calibration target is at this stage only an approximate delineation. Since the calibration target is used globally over all features of the image to counteract variations in instrument performance, it is critical that its detection algorithm is robust and precise. Errors perturb all measures on that image. Unfortunately, robust identification of calibration target segments is challenging as their pixel intensities often are very similar to those of pixels in the surrounding area. To solve this issue, the calibration target image section is first precisely automatically trimmed by Scan-o-matic.

###### 3.4.1.3.1 Orthogonal trimming of calibration target image

Due to the dimensional proportions of the calibration target (length much exceeds width) and the relative uniformity along its orthogonal axis, exact delineation orthogonal to calibration target axis is a less challenging task and is performed first (Fig S5A). In the orthogonal trimming step (along the axis that is perpendicular to the grey scale), pixel intensities across the calibration target image section are first normalized and then centred on zero. The image is then 2D convolved, using a 1D kernel, normalized and centred on zero and based on the target manufacturer’s target values and the expected length of calibration target segments. The vector containing the convolved image’s column maxima is smoothed using a 1D Gaussian filter. The standard deviation of the Gauss filter is set to half the width of a standard calibration target. The smoothing means that the signal peak will correspond to the calibration target centre, along the orthogonal axis. The boundaries of the calibration target strip are finally defined as being half the width of a standard calibration target away in each orthogonal direction. The image is trimmed at these boundaries.

###### 3.4.1.3.2 Parallel trimming of calibration target image

The parallel trimming starts with calculating the variance along the orthogonal axis (across the width of each pixel row). The outcome is convolved with a uniform kernel, with kernel size corresponding to the length of a standard calibration target. Permissible signals, a Boolean vector, are indices whose values do not exceed 1/40 of the signal range value. Limit is heuristically set. Potential calibration target edges are detected as non-zero outcomes of the [−1, 1] convolution of the permissible signal. The first edge is verified to have the sign that indicates a potential edge into the calibration target area, else all values in the candidate edges vector are shifted one to the right and the first image section index is inserted as first potential edge into a calibration target area at the first index of the vector. The last edge is then verified to have the sign indicating a potential edge out of the calibration target area – else the last index of the image section is appended as potential edge out of calibration target area. The true pair of edges that define the calibration strips boundaries in the vertical dimension is then selected as the pair of edges with an edge-edge distance that best agree with the expected length of the calibration target used. As pixel intensities of the brightest segment of the calibration target are virtually indistinguishable from the surrounding area (i.e. intensity difference ~ 0), the edge at the bright end of the calibration strip is not perfectly defined by this method. The brightest segment may therefore sometimes be abbreviated or extended. We buffer such abbreviations in the following way: from the centre position between the selected edges, we trim at half the expected length of the calibration target - plus an extra 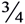 of the expected length of a segment of the used calibration strip - in each direction.

###### 3.4.1.3.3 Defining edges and centre points of calibration target sections

The trimmed area of the calibration target is further analysed along its vertical (along the sequence of segment areas) axis. First, the pixel intensity mean along the vertical axis is convolved with a [−1, 1] kernel. Signal spikes surpassing a heuristically set limit of 1.2 are candidate segment edges. These signals are trimmed so that only signal centres remain. The centre positions of signal spikes are aligned to the empirical expectation for spike distances (average distance between segment edges in the used calibration strip) such that the error is minimized. Any missing segment edges are filled in by interpolation. Exact distances between edges are calculated. Segment centre positions are interpolated from edge positions. Together with edge positions, these form a complete model of the calibration target area. The complete model is verified to fall within the trimmed image area.

###### 3.4.1.3.4 Pixel intensity calibration

Within each segment of the model of the calibration target, the most central 40% of pixels, in each dimension, are considered to be representative of pixels in the segment. The median pixel intensity of these pixels is further considered to be the most robust representation of pixel intensities in the segment. By excluding peripheral pixels and extreme pixel intensities, this provides stability in the face of physical damage to the calibration strip, in terms of scratches, dust and tears. The polynomial fit between segment pixel intensities and supplier provided calibration target values is determined, assuming a third degree polynomial function, *y* = *ax*^3^ ^+^ *bx^2^* ^+^ *cx*, where *x* = supplier provide target value (Fig S5B). All coefficients, *a, b* and *c* must be positive. This polynomial function is specific for each image and translates observed pixel intensities to a normalized value space of calibrated pixel intensities. Thus, it serves to make further analysis of pixel intensities independent of fluctuations in scanner properties over time and space and independent of variations between scanners.

##### 3.4.2 Image Analysis

Second pass analysis, called “analysis” in the user interface, detects, segments and quantifies the cell density of colonies as described in Fig. S2C. Analysis is performed in reverse chronological order, after the experiment has been completed, because detection and segmentation of colonies is more accurate on the later images when the cell density in each colony is maximal. As the second pass image analysis progresses from the last to the first image in the experimental series, analysis of each new image uses the candidate colony location, “blob”, from the previous iteration such that the two-dimensional mass centres are roughly maintained, i.e. colony expansion is assumed to have been more or less symmetrical in the horizontal plane.

###### 3.4.2.1 Grid placement

A virtual grid matching the user supplied pinning matrix is placed over each plate with the intention of aligning grid intersections with the centre of candidate colonies.

The process is described in Fig S6. First, each raw image of a plate (Fig S6A) is partitioned into sections by a randomly seeded Voronoi diagram. Each section is then Otsu-thresholded (Otsu 1975). For each section, the pixel intensities of background pixels form one Gaussian distribution whereas pixel intensities of colony pixels form another Gaussian distribution. Otsu-thresholding defines, for each pixel position, a threshold intensity value for that pixel to be considered a candidate colony (blob) pixel. The surface of thresholds, in each section, is heavily smoothed. Thresholds are then invoked to call blob pixels (Fig S6C). Sectioning is essential because background distributions of pixel intensities vary across a plate because of borders, inclusion of parts of the fixture, specks of dust, scratches etc. After applying the Otsu-threshold, an aggregation of blob pixels (referred to as a blob), are simplified and smoothed, using iterations of binary erode and binary propagation (Fig S6D). All blobs are evaluated for size (>40 pixels in area size, cover less pixels than the square of the expected distance between colonies) (Fig S6E), and for shape (expected to be circular due to the shape of the pin head that places the cells on the plate and expected to approximate evenly radiating growth; bounding box is expected to be roughly square) (Fig S6F). Thresholds for evaluation of size and shape are heuristically set based on observation of a large number of true (colonies) and false (hairs, dust specks, scratches, bubbles) blobs. Blobs failing to pass thresholds are discarded. The two-dimensional mass centre coordinates for retained blobs are stored in a NumPy array – an n-dimensional, indexed, uniform data structure. The average empirical spacing between pins, about 52 and 105 pixels in 1536 and 384 formats respectively, is used to construct an idealized grid. This idealized grid assumes that all pin centres, and consequently the centre of all deposited colonies, are an average distance apart. This idealized grid is fitted to the stored blob array such that the sum of errors between grid intersections in the idealized grid and the centre positions of the corresponding blobs are minimized. Grid intersections of the idealized grid is finally aligned with the nearest blob mass centre, given that the blob is close enough (squared distance must be less than a heuristically set threshold of 105 pixels).

###### 3.4.2.2 Colony detection, segmentation and pixel intensity analysis

For each grid intersection, corresponding to a pinning location, the surrounding local area is assumed to describe a mixed Gaussian model, with blob and background pixels distributed in two Gaussian distributions (Fig S7A, B). Each local area of the raw image is first smoothed with a median filter (median filter; size 3×3 pixels) to reduce noise (Fig S7C). The local area is then thresholded with an Otsu-threshold, as in 3.4.2.1, to distinguish candidate colony/blob pixels from background pixels. To ensure that the edge of each colony is included in the colony’s definition, blobs are passed through iterations of binary dilation, also as in 3.4.2.1. At this stage, blobs falling inside, or partially inside, other blobs, are merged. The largest, most circular blob and which, if it’s not the first image to be analysed, best concurs with the blob of the previous iteration (image, analysed from end to start), is selected. Other blobs are placed in a second array as trashed pixels. Trashed pixels correspond to specks of dust, scratches on the plastic, or other similar types of noise and are not considered further. The local background is then defined as the compliment of the union of the blob and the trash arrays, with an eroded safety margin (Fig S7C). The mean of the interquartile range (IQR) of the local background is used as a scalar measure of the transparency of the solid medium in the local context of the colony considered. The IQR mean is not as vulnerable to outlier values stemming from the noise discussed above as mean, and is more precise than a median of discrete values, such as image pixel values. The difference between the pixel values of the blob and the local background value represents the per pixel darkening effect of the cells in the corresponding area of the plate. These values are retained for downstream use.

###### 3.4.2.3 Converting pixel intensity to cell density

The transformation from cell darkening values to actual cell densities is based on a calibration experiment in which 42 colonies with a wide range of population sizes were analysed in Scan-o-matic, according to the procedures described above. Solid media (agar) pieces with each colony on top were then carefully removed from the plate, colonies were washed off from the agar into sterile water by extensive vortexing and the actual number of cells in each colony was estimated using two independent techniques. First, OD600 in a spectrometer (Pharmacia Biotech NovaspecII) was measured and transformed into cell density based on that 1 mL of OD = 1.00 medium corresponds to 10^7^ cells. Second, cells were sonicated and counted using a Fluorescence Activated Sorting Machine (FACS) (FACS; BD FACSAria). The two cell density measures (OD and FACS) showed close to perfect linear correlation (Fig. S8A), ensuring that they accurately capture, or at least scales linearly with, real cell numbers in each colony. OD-measures were therefore used to transform pixel darkening values (calibrated pixel intensity) to cell density, assuming a polynomial relation (Fig S8B, C). The best fit polynomial was used: *y* = 3.3797963108805451*10^−5^\**x*^5^ ^+^ 48.990614276885069*x*, for *x* > 0, where *y* = cell density and *x* = pixel darkening values. Intermediate exponents are omitted to avoid over-fitting the curve. Pixels with *x* < 0, are set to *x* = 0 before applying the polynomial. The shape of the calibration polynomial curve indicates the range for which the scanner manages to reliably resolve the number of cells per pixel. With extremely strong pixel darkening effects, transformation to cell density space is not reliable. Thus, a maximum capping threshold is enforced such that for *y* > 2500, *y* is set to 2500. If capping is invoked, a warning is emitted. Note that the threshold is much beyond what is observed for white-coloured yeast colonies. However, if colonies have a very dark coloration, due e.g. to intracellular accumulation of metals such as tellurium (Ottosson et al. 2010), the capping may be invoked. We strongly caution against scanning on media inducing such dark coloration and independent calibration experiments must in these circumstances be performed to obtain alternative polynomial fit coefficients. Cell count estimates per colony and time are stored in NumPy files and XML-format.

### 4. Extracting population growth parameters from dense growth curves

#### 4.1. Smoothing of growth curves

To extract growth features from growth curves, cell count estimates are first re-arranged to a per colony time series, rather than per scan, providing growth curves. A smoothing function is then applied to each curve. The smoothing consists of two parts. First, a median filter of size = 5 (i.e., the median of each five consecutive OD values replaces the third value in each such series of five) is employed. Reflect edge conditions allow filtering of initial and end measurements. The median filter removes individual noisy estimates (positive and negative spikes) but does not smooth data beyond that. Second a Gaussian filter of width *σ* = 1.5 is employed. In essence, this adjusts each estimate based on all other estimates in the curve with each estimate given adjustment weight depending on how far (in time) it is from the estimate to be adjusted. Weights follow a normal distribution centred on the estimate itself, with the standard deviation of this weight distribution set to 1.5. From the smooth curve a range of phenotypes are extracted. A phenotype in this context is any quantitative feature of the growth curve. Our focus in this work is to generate high quality measures of population doubling time. Auxiliary phenotypes relevant to this aim are: initial population size (for correlation studies), error of linear regression at the time of maximum growth (for quality control), time of maximum growth (for quality control) and the fit of an extended Chapman-Richards model (for quality control).

#### 4.2 Extracting population size doubling time, error of the linear regression at maximum growth rate and time of maximum growth rate

We assume the budding yeast life-cycle on solid medium to be completely mitotic (i.e. no meiosis and no hyphae formation is assumed to occur), the length of the mitotic cell cycle to be normally distributed and cell death to be negligible. In these circumstances, a population is expected to double in size during the time of an average cell cycle. Consequently, a growth-curve with a log(2) y-axis should be linear as long as environmental conditions are constant from the perspective of the cells. In reality, both the genetic and the non-genetic environment of each cell is in constant flux and assumption of an extended phase of such linear growth on a log(2) scale is clearly precarious. Growth is rarely log(2) linear for any extended period of time, and this becomes increasingly evident when growth is precisely estimated at short intervals. We solve this by brute force. To reliably measure the maximum slope of the smoothed growth curve we do all possible linear regressions with a window size of five. From these we select the linear regression with the steepest slope and calculate the doubling time and the error of the regression and record these. The time for the centre point of the curve segment is stored as the auxiliary phenotype time of maximum growth.

#### 4.3 Initial population size

To obtain a robust measure of the number of initially deposited cells, we calculate the mean of the initial three (from the first three time-points) population sizes.

#### 4.4 Extended Chapman-Richards Four Parameter Curve

We extended the classical Chapman-Richards four parameter model of a growth curve use it to obtain a measure of the reliability of calculated generation times. The model is not used for extracting the generation times themselves, only to indicate growth curves which are of potentially lower quality and for which generation times should be considered with caution. The extension corresponds to inclusion of an initial y-offset, *D*, which makes the model better fit to our data as the curves at *t* = 0 tend to start around 2^15^ cells rather than at the classical Chapman Richards assumption of *y* = 0. The model fit is done in log(2) y-value space. To ascertain that parameters are kept within their defined ranges, the input parameters to the model are transformed as follows:

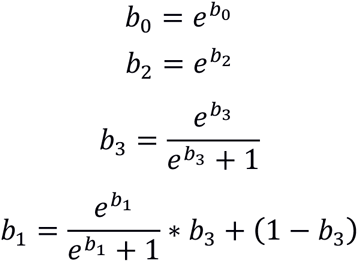

Initial parameter estimation was optimized iteratively with parameter estimates for the first iteration selected being based on previously suggested estimates (Pylvänäinen 2005). In consecutive iterations, the mean parameter values from the previous iteration, for over 6,000 curves, were used as new parameter estimates. Values producing the least summed error of fit and used by Scan-o-matic were: *b*_0_ *=* 1.64, *b*_1_ = −0.1, *b*_2_ = −2.46, *b*_3_ = 0.1, *D* = 15.18. The model fit to each growth curve was recorded and used as an approximate quality measure of that curve, i.e. a poor fit was taken to indicate a growth curve that is suspicious and in need of further attention.

### 5. Quality Control and Normalization of Population Growth Data

#### 5.1. Quality control

To give a visual representation of the distribution of phenotypes over a plate, we designed a user interface that shows each plate as a two-dimensional heatmap. The coordinates reflect colony positions and the colour intensity reflect the phenotype value. The phenotype to display is selected from a drop-down menu. In the lower part of the display, the currently selected colony’s growth curve is displayed together with phenotype information. The plate selector allows toggling between the four plates of the fixture, one at a time. The phenotype drop-down selector allows selection of which phenotype to display in the heatmap format. The key output phenotype is the generation time/doubling time). Other phenotypes are: “Error of GT”, i.e. the error of the linear regression at the time of the population size doubling time extraction, the “Fit of the modified Chapman-Richards model” and “time at maximum growth rate”. A large error of the linear regression, a poor model fit and a very early or very late extraction of population size doubling times suggest growth curves of low quality. Such suspected low quality growth curves can and should be manually inspected by selecting individual positions on the heatmap. Scan-o-matic will automatically display the colonies in ascending quality order to ease inspection. Low quality growth curves should be discarded before normalization, not to influence the normalization procedure. We urge users to be conservative in retaining growth curves, in particular if replication on plates is high. In the reported experiments ~0.3% of curves were excluded. During quality control, meta-data for the experiments can be joined with the observed phenotypes by importing a suitable spreadsheet file. This establishes a connection between the growth curve inspected and a description of the colony. The spreadsheet file needs to meet these requirements:

1. Each row on a sheet represents all information about a position on the plate.
2. Each row on a sheet only represents data for one position on the plate.
3. Order of rows in a sheet should correspond to first left to right order of colonies in a plate row and then a top to bottom order of plate rows (assuming a landscape orientation of each plate
4. Each sheet represents a complete plate, a fourth of a plate or a collection of several full plates.

Both OpenOffice and Microsoft Excel file formats are supported. Examples are provided <u>here</u>. The use of four separate sheets with a quarter of the number of strains needed to fill the intended experimental plate in question (e.g. 4 sheets with 384 rows for a plate with 1536 positions) will automatically trigger Scan-o-matic to interpret them as partial plates to be interleaved. For each of the four, the information will be placed on every second column and every second row. The positions of the first sheet will have no offset to its origin. The origin of the second is offset along the long axis, third along short and fourth along both. A schematic of the interleaving is shown in Fig. S12.

#### 5.2 Normalization

A fundamental challenge is accounting for systematic local variations within a single plate. Such systematic bias creates unacceptably high rates of false positives and negatives, even when measurement precision is excellent. To overcome such local variations, we introduced a dense matrix of genetically identical controls, sacrificing every 4^th^ experimental position and replacing it with a control (Fig S11). We accept the corresponding small reduction in throughput. The growth of these controls is used to establish a three-dimensional surface of generation times that well captures local environmental variations affecting growth. The three-dimensional surface is established in three steps. First, controls with extreme values are removed before actual normalization, to avoid distorting the normalization surface. This is done iteratively by excluding the 4% of controls whose phenotype values are >2σ away from the mean of controls, given a normal distribution assumption. Filtering is done iteratively <10x, recalculating mean and standard deviation after each removal until on average ten controls have been removed. Admittedly, this filtering does remove some useful curves from the building of the reference surface. However, the exclusion of truly dubious positions that otherwise would severely distort the normalization surface far outweighs this loss. Second, using all remaining control positions, the normalization surface is extended to cover all other positions (non-control and removed control positions). This is done by splining/interpolation according to the method “griddata” of SciPy’s “interpolate module”. This is a cubic splining, i.e. all nearest neighbours, vertically and horizontally, are used for the interpolation. When a non-control position has been filled, it in turn serves as a point of departure for filling in neighbouring unfilled positions. This is particularly important towards edges, where controls are sparser. In essence, the interpolation procedure provides a rough estimate of the expected generation time of all positions, assuming that they had been controls. Third, this reference-grid normalization surface is smoothed in two steps. First, a median filter with a 2D kernel of size 3 × 3 is applied to remove noisy, i.e. individual points deviating from their surroundings. The median filter was chosen because it preserves the sharpness of larger features. Second, a Gaussian filter with size σ = 1 is applied. In essence, this adjusts the value in each position by taking the value of surrounding positions into account, with the contribution of each neighbouring position being weighted by its distance to the position of interest. The Gaussian filter vastly improves the rather crude estimates of colonies along the two edges on which no control colonies exist. When the surface has been completed, the log(2) difference between the actually observed population size doubling time for a position, and the value of the normalization surface in that position is calculated. This relative growth measure, which estimates the performance of a colony in relation to what a control would have had – if it had been in that position - is reported as the final output. For simplicity, we refer to it as the log(2) relative population size doubling time.

### 6. Data Publication

Both final and raw phenotypes can be exported in a tab separated format. Phenotype values are appended after the user given meta-data for each colony/row. If no meta-data is given during quality control, phenotypes are preceded only by the positional information of the experiment. Raw growth curves are stored as NumPy files, one per scanned image. Each file is organized according to plate, row, and then column. The raw growth data is also stored in an XML-format (either as full tag names or shortened aliases). Specifications for the XML are available in the Scan-o-matic repository documentation.

### 7. Access to Scan-o-matic

Scan-o-matic is currently available as two separate packages. 1) Scan-o-matic for scanning and analysis is available here, including a demonstration project. 2) Scan-o-matic for quality control and normalization is available here. 3) https://github.com/local-minimum/scan-o-matic provides up-to-date information on new versions of the program, access to a wiki, a help section and opportunity to report errors.

### 8. Wet-lab experimental procedure

Solid media cast in plates designed for use with Singer RoToR HDA robot (Singer Ltd) was used. Each plate was cast with precisely 50mL of Synthetic Complete (SC) in the form of 0.14% Yeast Nitrogen Base, 0.5% ammonium sulphate, 0.077% Complete Supplement Mixture (CSM, ForMedium), 2% (w/v) glucose and pH buffered to 5.8 with 1% (w/v) succinic acid. Media was supplemented with 20g/L of agar. Casting was done on an absolutely level surface with drying for ~1 day. Where indicated, 2% glucose (unstressed environment) was replaced by 2% galactose and/or supplemented with 0.85M NaCl. Two strain layouts were used: 1) all colonies being diploid BY4743 reference strain (Brachmann et al. 1998) (Fig 1–3) 2) colonies corresponding to single yeast gene knockouts of the haploid BY4741 deletion collection (Giaever et al. 2002), with BY4741 control colonies interleaved in every fourth position (Fig 4). In the first case, a lawn of perfectly mixed BY4743 cells was cast by dispersion of 50 μL of an overnight culture evenly across the surface of the solid agar media and incubation at 30°C for 2 days. From this lawn, a 1536 colony format pre-culture (incubated for 2 days) was initiated using the RoToR pinning robot and the short pin 1536 pinning pad. In the deletion strain experiments, frozen glycerol stocks were first pinned using 96 format long pin-heads into 96 well microplates containing liquid SC media. These were cultivated for 2 days before being transferred to 384 solid SC media (with the use of 96 format long pin-heads), and incubated as pre-pre-cultures (2 days incubation). In parallel, a lawn of BY4743 control strains was constructed as above, pinned onto 384 solid media and allowed to grow for 2 days. Deletion strains and controls strains were interleaved in a 1536 format (with 384 short-pin pinning pad) using a custom-made pinning program for the RoToR robot as described in Fig S11. Plates were pre-cultured for 2 days, and then re-pinned using short 1536 pins to initiate actual experiments. Cultivations were performed at 30°C with n=3 replicates of each strain in juxtaposition, as a consequence of the pinning scheme (Fig S11). For the confirmation experiment (Fig 4E-F, S12B-C) the same procedure was employed, except that n=24 replicates were used, distributed over two experimental plates. For the reference liquid media experiments (Fig 4E-F, S12B-C), 10μl from each well from the same 96 format microtitre plates that were used for the agar experiment was transferred to 350μl SC media per well in a 100-well format Bioscreen C plate. Pre-cultures were incubated for 2 days at 30°C. 10μl of each pre-culture was then transferred to six experimental plates, each well containing 350μl media (media as above but excluding the agar) and incubated in six different Bioscreens (n=6) for 3 days, as previously described (Warringer et al. 2011).

## SUPPLEMENTARY FIGURE LEGENDS

**Fig S1.**
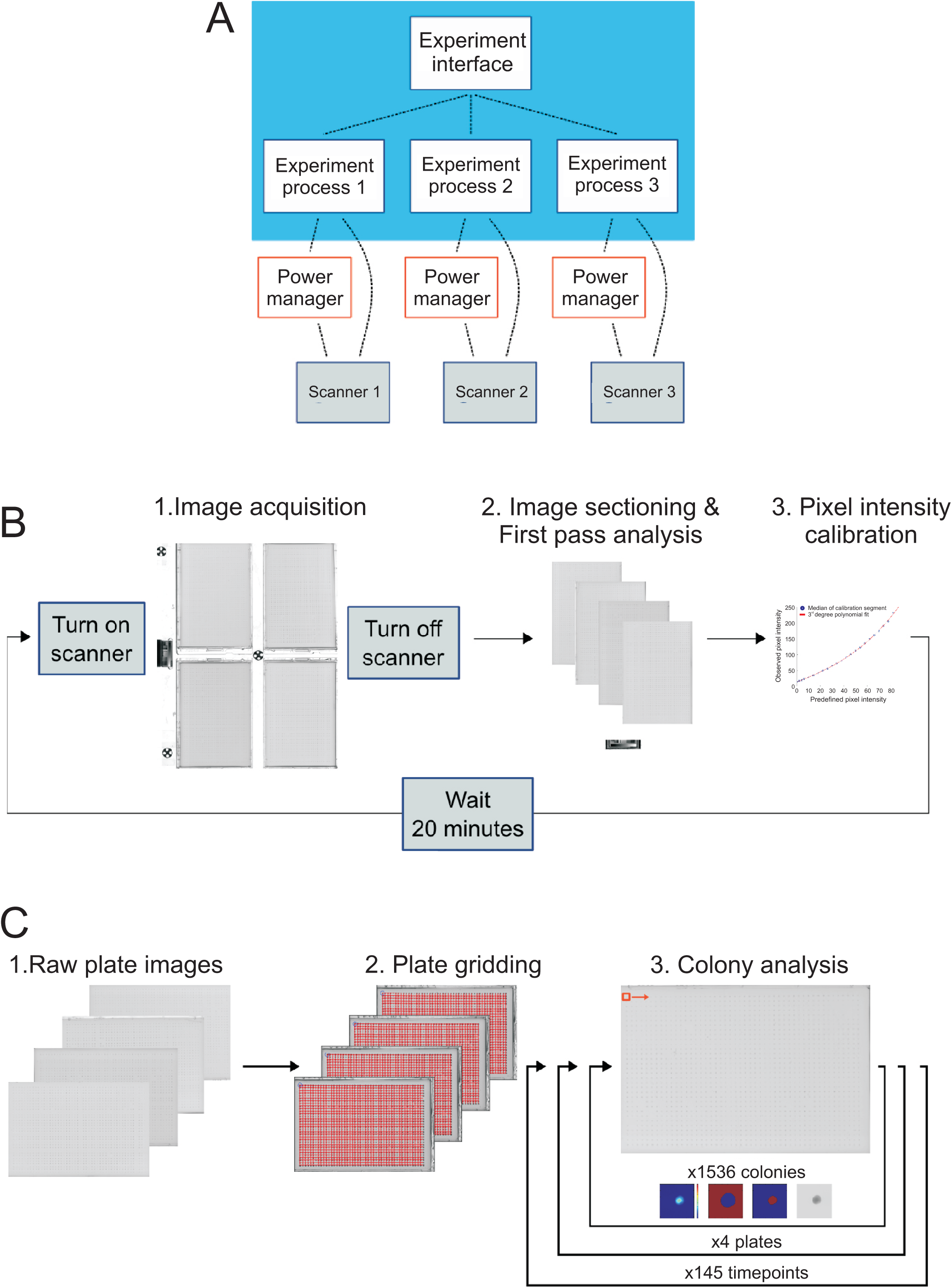
Scan-o-matic process overviews. A) Overview of the Scan-o-matic hardware – software arrangement and interaction. Three Epson Perfection V700 PHOTO scanners (Epson Corporation, UK) are connected via USB to each controlling computer. The power supply of each scanner is controlled individually and independently from the computer using a single GEMBIRD EnerGenie PowerManager LAN with multiple sockets (Gembird Ltd, the Netherlands). Software structure is shown inside the blue rectangle. Experiments on the different scanners connected to one computer are run as separate autonomous processes, but scanner power supply toggling is coordinated by the server. This ensures that malfunction in one scanner does not affect other scanners connected to the same computer. B) Overview of the first pass analysis. The first pass analysis identifies the position of orientation markers within each image and matches recorded positions to positions in a stored model of the used fixture (see Fig S2B and S4A). The match is used to precisely determine positions of each of the four plates and of the transmissive scale calibration strip in the respective image (see Fig S3B). These are sectioned out and stored as separate image features. The transmissive scale is further analysed to establish positions of transmissive scale areas (Fig S4A) and used to calibrate pixel intensities so that these become independent of variations in scanner properties over time, space and scanners (Fig S4B). Observe that the first pass analysis is performed as images are acquired, image per image, and is completed, reported and stored rapidly after acquisition of each image. The experiment processes rests until next scan is expected. C) Overview of second pass analysis. Second pass analysis detects colonies and quantifies their cell density, in each image. Second pass analysis is performed in a time reverse manner, i.e. starting from the last image in each time series, after all images have been acquired. First, a grid corresponding to the pinning density used is placed on top of each plate image and positions of grid intersections are precisely aligned with the centre point of candidate colonies (Fig S5). Second, each grid intersection region is segmented so that colonies are distinguished from background – pixels are assigned to colonies and local backgrounds respectively (Fig S6). Pixel intensity differences between the pixels of each colony and its local background average pixel value are converted into cell density by calibration to empirical measures (Fig S7). The complete process is iterated for all 1536 colonies, in each of the four plates, for each of the images (145 for a 48h experiment with 20 min measurement intervals), starting from the last image.

**Fig S2.**
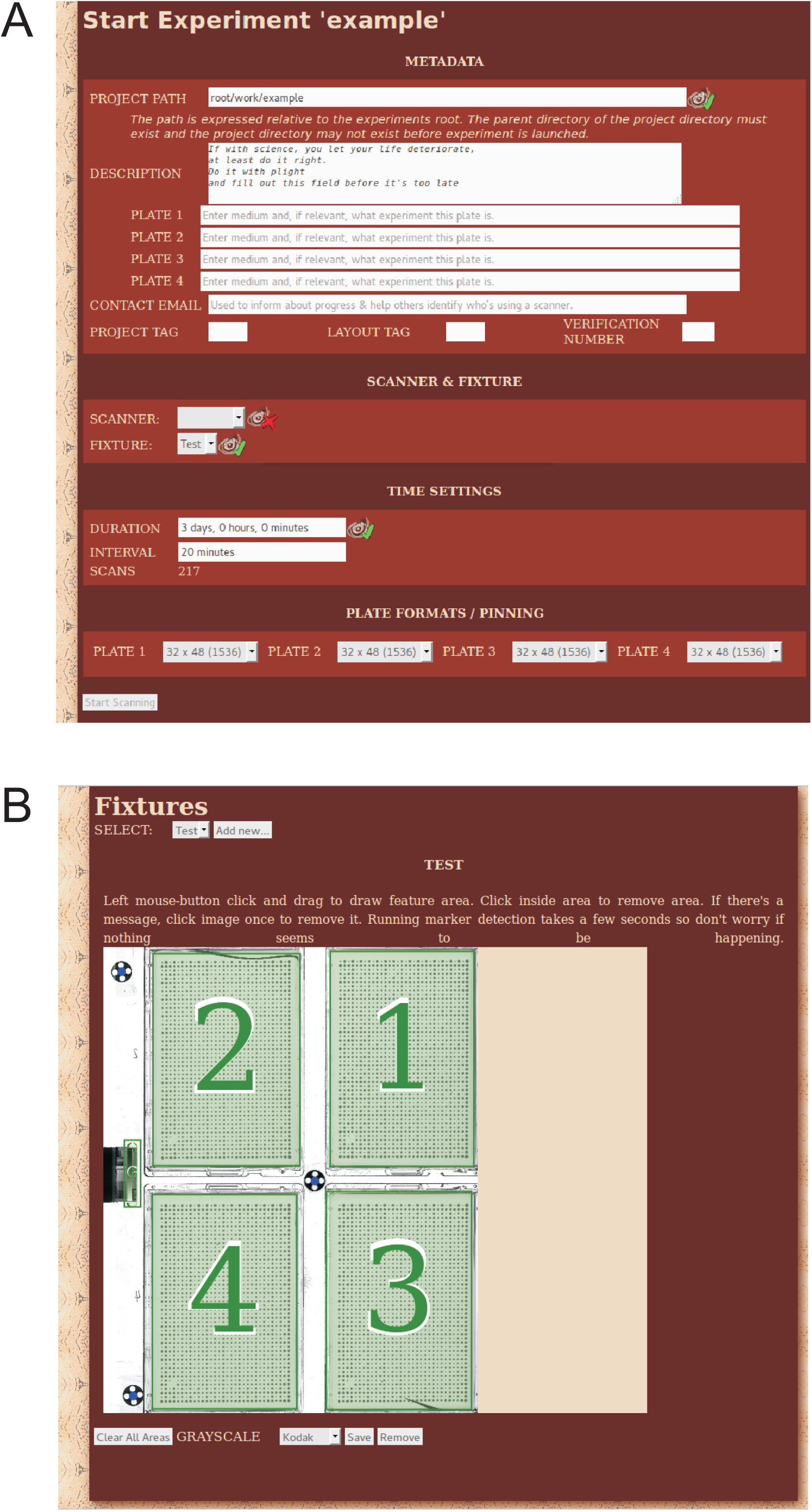
Pre-experimental Scan-o-matic user interfaces. A) User interface for initiating a Scan-o-matic experiment. Parameters that are set are: experiment duration (from 14 minutes to 7 days, but typically 2–3 days), time interval between consecutive scans (from 7 to 180 minutes but typically 20 minutes) and the pinning density of each plate (96, 384, 1536, 6144). The four plates in each scanner will by necessity be subject to the same time and interval settings. The experiment is initiated from the interface and will run until completion. B) Scan-o-matic fixture calibration user interface. Before experiment, once per scanner-fixture combination, a spatial calibration model that describes the layout of the fixture needs to be created using this interface. Scan-o-matic uses this fixture calibration model to section images of said fixture into their meaningful components. The fixture name and the number of fixture orientation markers are given. Plate positions, and the position of the transmissive scale calibration target strip are marked by interactive click-and-drag mouse maneuvers. The type of calibration target is selectable from a drop down menu.

**Fig S3.**
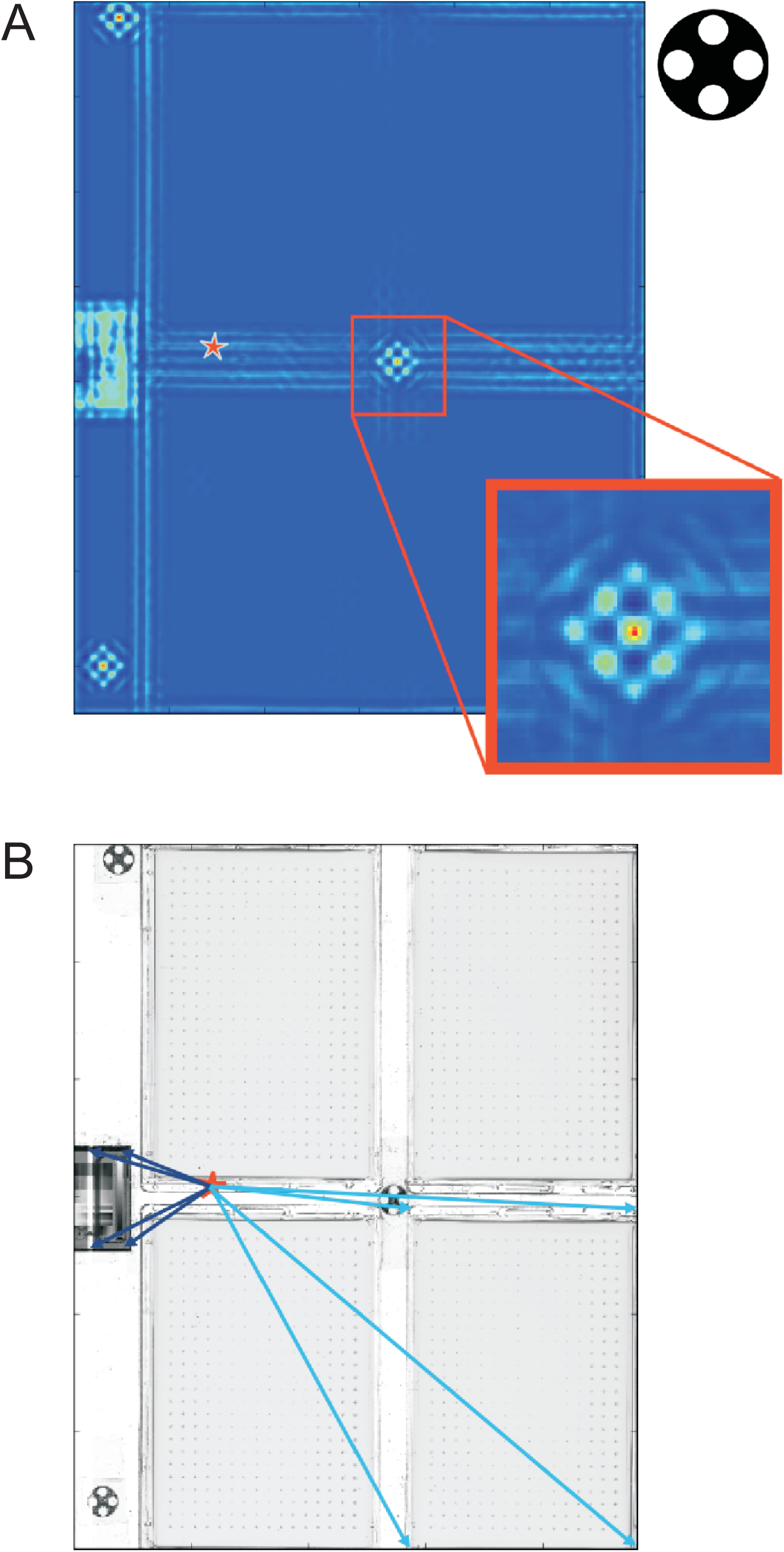
Localizing and positioning image orientation markers relative the fixture calibration model. A) Localizing orientation markers within each individual image. Each image, I, is thresholded with a pixel value of 127 (value range midpoint) resulting in a two colour image (dark and light blue). The thresholded image is convolved, using a fast Fourier transformation, with a black and white representation of a fixture orientation marker (upper right corner). This evaluates, for each pixel of the image, how well the local context of that pixel matches a representation of a fixture orientation marker. Iteratively, with the number of iterations corresponding to the number of fixture orientation markers in the fixture model, the strongest signal position is selected and chosen as the centre position (red) of that fixture orientation marker, *c. c* is stored as a marker coordinate. A strong signal is obtained when the majority of the adjacent pixels correspond to the expected marker two-colour pattern. To avoid repeated reporting of the same marker the local area, corresponding to the size of the convolution kernel, is set to one less than the minimum of the convolution surface. The process of marker localization is repeated on the modified surface until all markers have been found. The mass centre of all three markers (red star) is calculated to serve as a fixed reference point. B) Localizing and sectioning out plates and transmissive scale calibration strips. Orientation marker coordinates are matched between image and fixture calibration model and the offset vector between the two-dimensional centres of mass of the two coordinate systems (image and fixture calibration model) obtained. The image’s instance of the fixture model is adjusted with this offset vector so that a near perfect match between image and model sections is obtained. Each relevant image feature (plates and calibration target areas) is represented by four vectors (dark blue and light blue arrows) from the fixed reference point (two-dimensional mass centre, red star). Each relevant image feature is sectioned out and stored separately for later analysis (see Fig S4–8)

**Fig S4.**
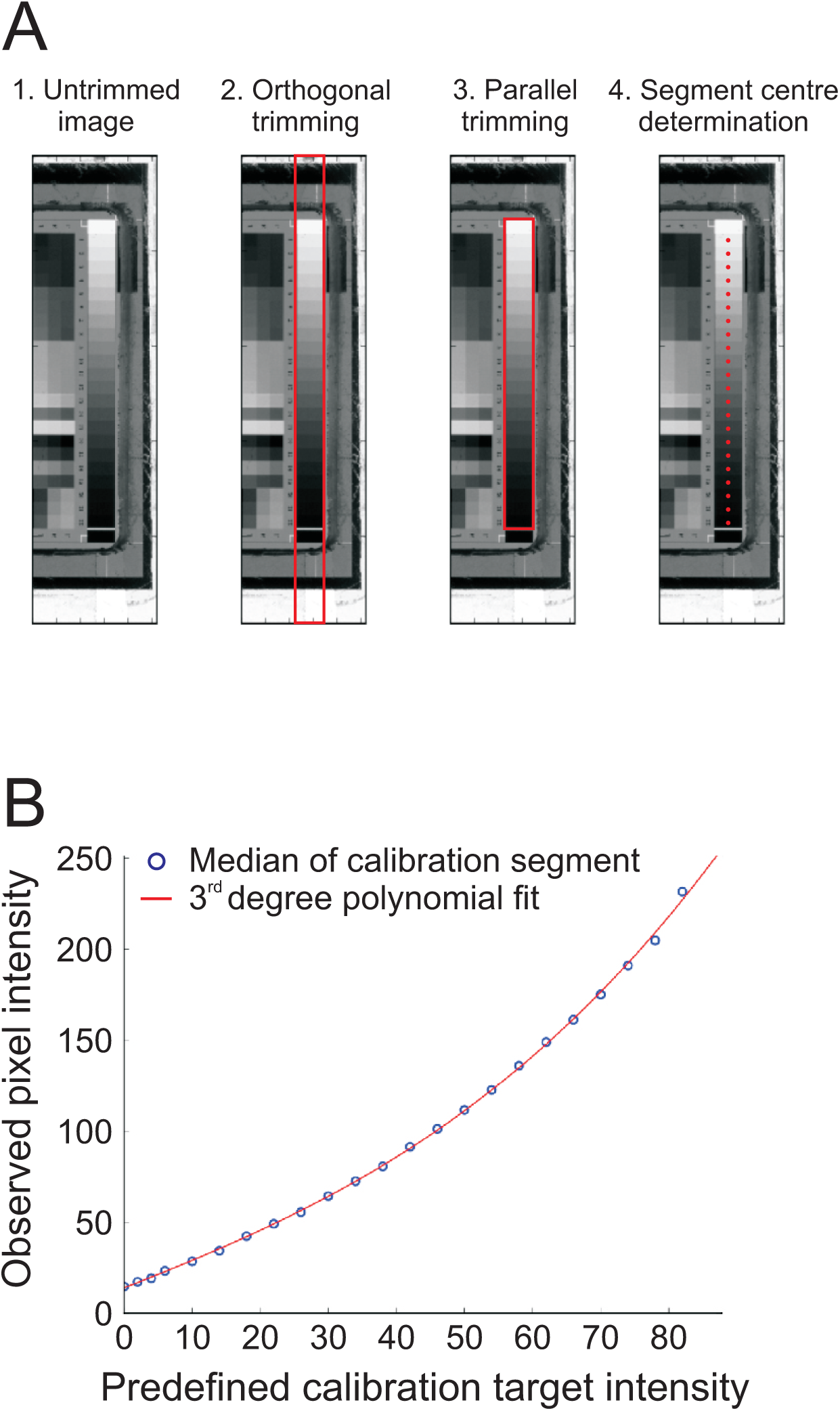
Calibrating pixel intensities using a transmissive scale calibration target strip. A) The stored image section corresponding to a transmissive scale is sequentially trimmed, first in an orthogonal and then in a parallel fashion, to exclude all pixels not belonging to a transmissive scale proper (red rectangle). Transmissive scale segment centres are localized (red circles). The median of the pixel intensities around each segment centre is taken as the best representation of pixel intensities in that segment. B) Supplier provided calibration target values for each segment is available (x-axis). Together with the detected transmissive scale segments’ pixel intensities (y-axis), these are used to construct a third degree polynomial function that is specific for each transmissive scale calibration target strip and image. The polynomial translates observed pixel intensities to a normalized value space of calibrated pixel intensities. These are used for all further analysis (see Fig S5).

**Fig S5.**
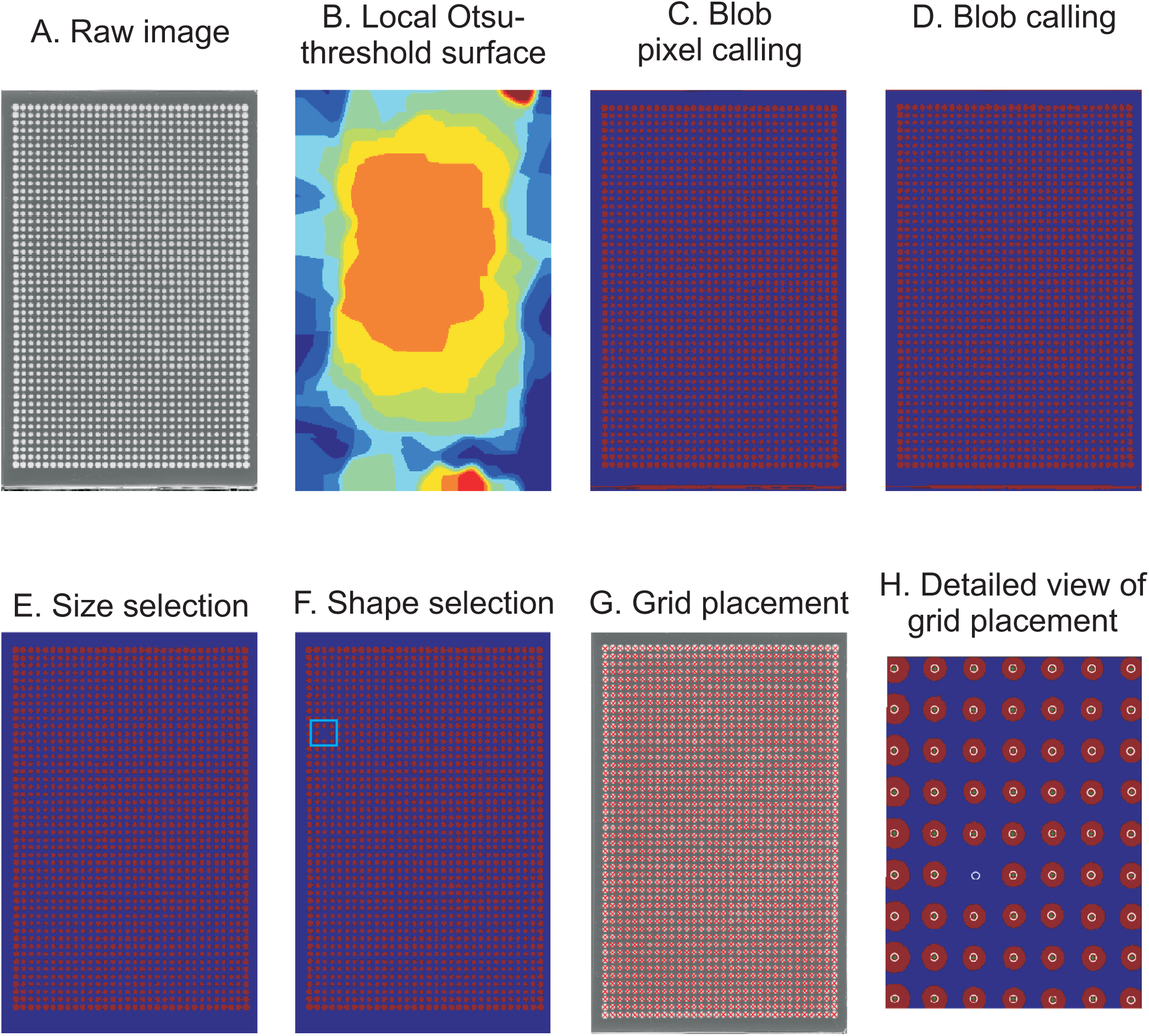
Placing a virtual positioning grid across each plate image. The first step in the second pass analysis places a virtual grid that matches the user supplied pinning matrix over the raw image of each plate and aligns grid intersections with the centres of candidate colonies. A) Raw image of a plate. B) Otsu-thresholded surface of a plate section that corresponds to a zoom-in on a candidate colony on the plate in A. Each image is first partitioned into sections by a randomly seeded Voronoi diagram. Each section is then Otsu-thresholded. For each of these sections, the pixel intensities of background pixels form one Gaussian distribution and the pixel intensities of colony pixels form another Gaussian distribution. The Otsu-threshold defines the optimal intensity for each mixed Gaussian distribution to separate the component distributions into pixels considered as candidate colony (blob) pixels and background pixels. For all the sections in the plate, the surface of threshold-values is heavily smoothed. C) Thresholds are invoked to call blob pixels (red) as distinct from local background pixels (blue). Note the thin red line of false positives at the bottom of the image. D) After applying the Otsu-threshold, aggregations of blob pixels are simplified and smoothed, using iterations of binary erode and binary propagation. This re-assigns some pixels at the blob/background boundary. E) All blobs are evaluated for size (>40 pixels in area size and cover fewer pixels than the square of the expected distance between colonies). Note that this removes the thin line of false positives at the bottom of the image. F) All remaining blobs are evaluated for shape (expected to be circular due to the shape of the pin head that places the cells on the plate and expected to approximate evenly radiating growth; bounding box is expected to be roughly square). Note the removal (blue rectangle) of a blob that could not unambiguously be established as a true colony. G) The average empirical spacing between pins in the used pin format is used to construct an idealized grid. This idealized grid assumes that all pin centres, and consequently the centre of all deposited colonies, are an average distance apart. The idealized grid is fitted to the stored blob array such that the sum of errors between grid intersections in the idealized grid and the centre positions is minimized. Grid intersections of the idealized grid are finally aligned with the nearest blob mass centre, given that the nearest blob is close enough (squared distance must be less than a heuristically set threshold of 105 pixels). H) Resolved view of a placed grid. Grid intersections (white circles) and blob centres (light blue crosses) are indicated.

**Fig S6.**
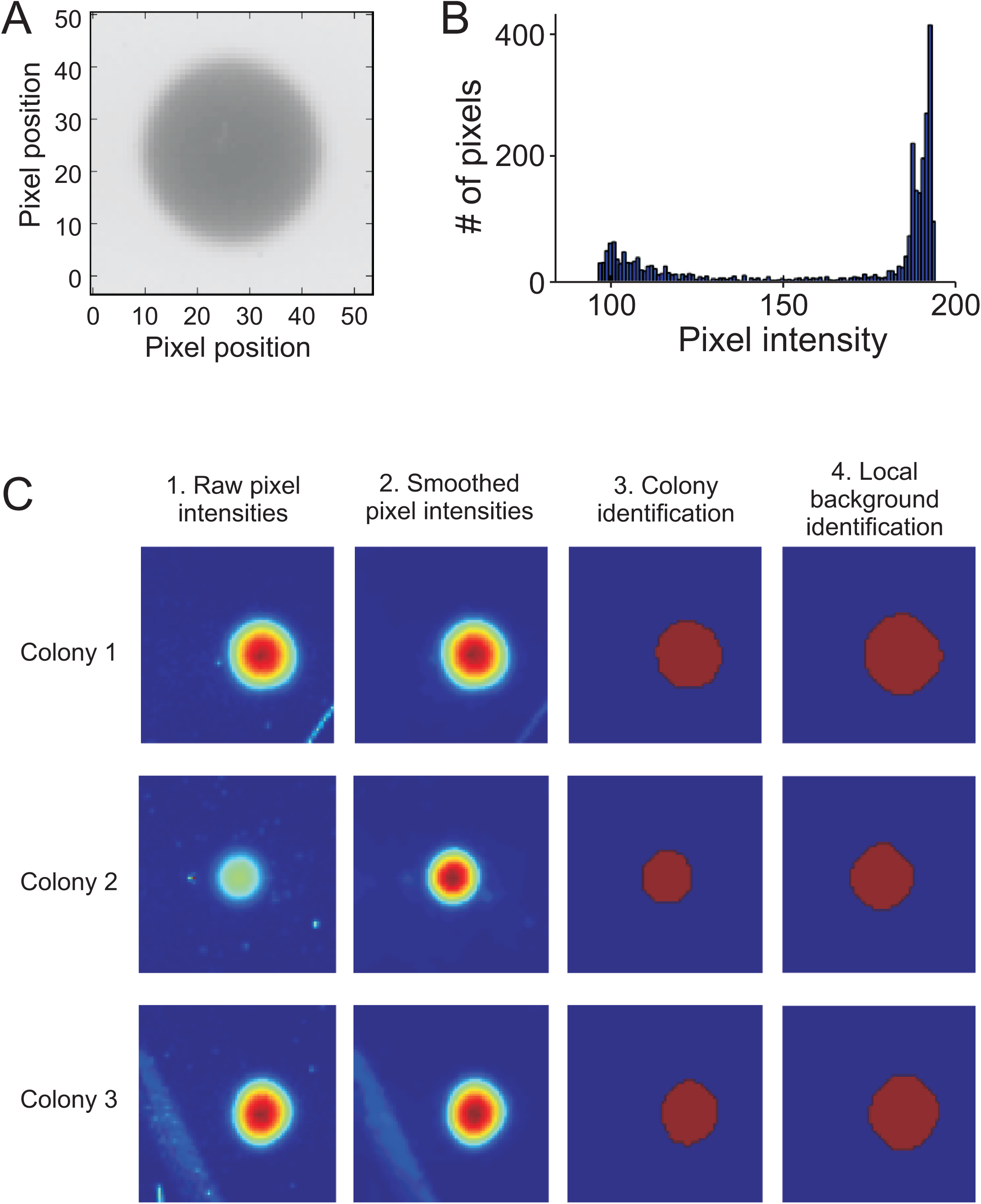
Estimating pixel intensities in each colony. A) Raw image of a random colony and its local background. B) Distribution of pixel intensities at the grid intersection shown in A. Scan-o-matic assumes that colony pixel intensities and local background pixel intensities follow two overlapping distributions. Pixels are assigned to colony and local background respectively based on their pixel intensity, as described below. C) Pixel intensities in three local areas (centred on three grid intersections) of an image. Dark red = high intensity, dark blue = low intensity. Areas reflect three complex challenges: hair (top panels), specks of dust (centre panels), and distortions in the transparency of the plastic casting (lower panel). The landscape of raw pixel intensities (far left panels) is first smoothed with a median filter (size 3×3 pixels) to reduce noise (left panels). Note that this vastly but not completely, removes the impacts of the challenging elements. Smoothed pixel intensities are then thresholded with an Otsu-threshold, i.e. assigned as candidate colony/blob pixels if they pass a local threshold. To accurately define blobs and ensure that the edge of each colony is included in the colony’s definition, blobs are passed through iterations of binary dilation. This re-assigns some pixels at the boundary of each blob. Blobs falling inside, or partially inside, other blobs, are merged. The largest, most circular blob that, if it’s not the first image to be analysed, best concurs with the blob of the previous iteration (image, analysed from end to start), is designated as a colony. Other blobs are placed in a second array as trashed pixels. Trashed pixels correspond to specks of dust, scratches on the plastic, or other similar types of noise and are not considered further. Note that this completely removes any remaining influence of the challenging elements encountered (right panels, blob pixels in red). The local background (far right panels, background pixels in blue) is then defined as the complement of the union of the blob and the trash arrays, with an eroded safety margin. The mean of the interquartile range (IQR) of the local background is used as a scalar measure of the transparency of the solid medium. The difference between the pixel values of the blob and the local background value represents the per pixel darkening effect of the cells in the corresponding area of the plate. These values are retained for downstream use (see Fig S7).

**Fig S7.**
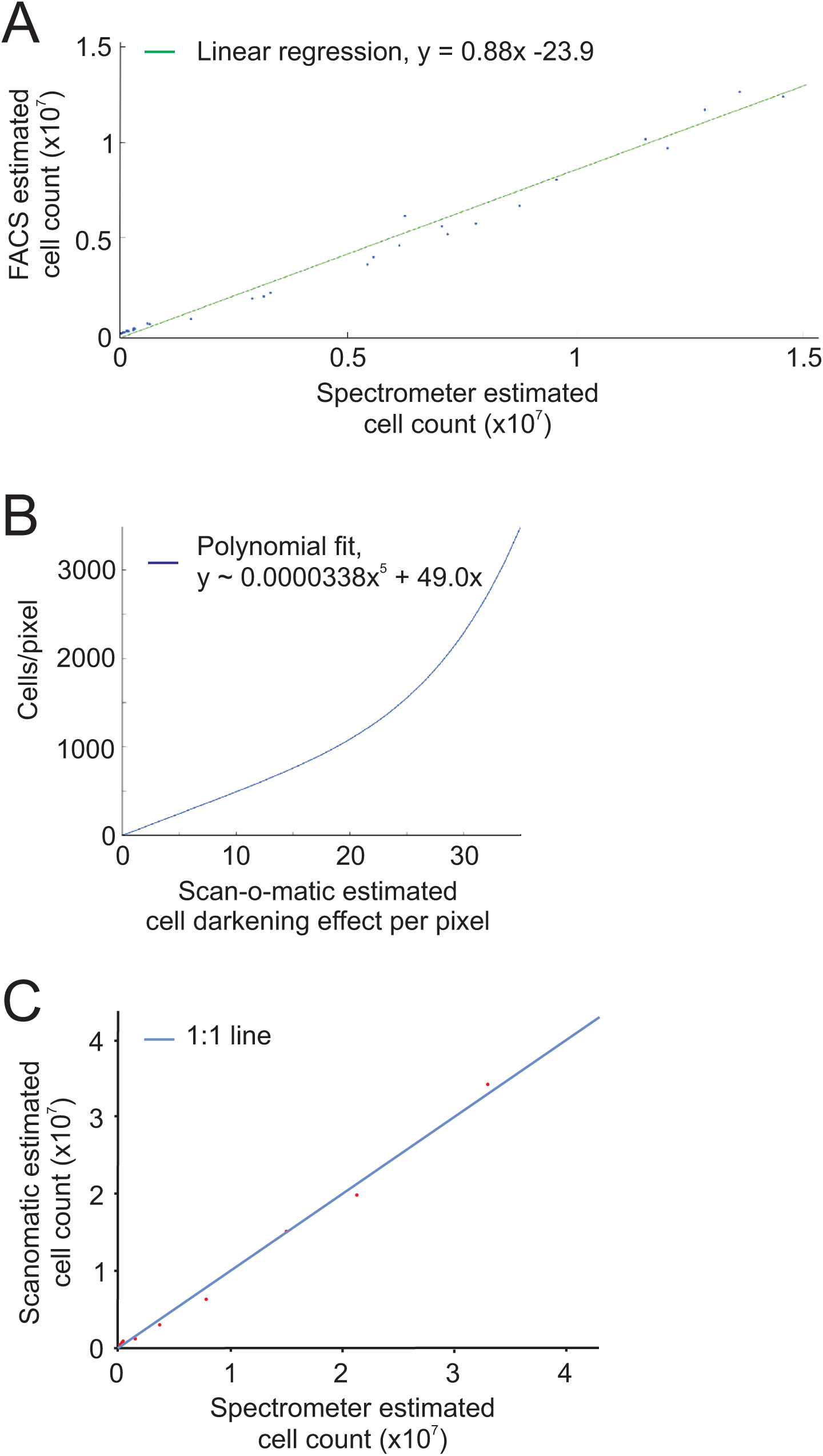
Converting pixel intensity to cell density. The transformation from cell darkening values to actual cell densities is based on a calibration experiment in which 42 colonies with a wide range of sizes were analysed in Scan-o-matic, according to the procedures described above. Solid media (agar) pieces with each colony on top were then carefully removed from the plate, colonies were washed off from the agar into sterile water by extensive vortexing and the actual number of cells in each colony was estimated using two independent techniques. First, OD600 in a spectrometer (Pharmacia Biotech NovaspecII) was measured and transformed into cell density based on that 1 mL of OD = 1.00 medium corresponds to 10^7^ cells. Second, cells were sonicated and counted using a Fluorescence Activated Sorting Machine (FACS) (FACS; BD FACSAria). A) The two empirical cell density measures (OD and FACS) showed close to perfect linear correlation, ensuring that they accurately capture, or at least scales linearly with, real cell numbers in each colony. B) OD-measures were used to transform pixel darkening values (calibrated pixel intensity) to cell density, assuming a polynomial relation. The best fit polynomial was used: *y* = 3.38*10^−5^\**x*^5^ ^+^ 49.0*x* (coefficients rounded for display purpose; higher precision was used in actual computation), for *x* > 0, where *y* = cell density and *x* = pixel darkening values. Intermediate exponents are omitted to avoid over-fitting the curve. Pixels with *x* < 0, are set to *x* = 0 before applying the polynomial. C) To verify the accuracy of polynomial transformation as a concept, 32 of the 42 colonies were randomly selected and a new polynomial constructed from them. The remaining 10 colonies were used to estimate the cell density as explained above and compared to the measured densities using the new polynomial. Close to perfect 1:1 correlation was observed.

**Fig S8.**
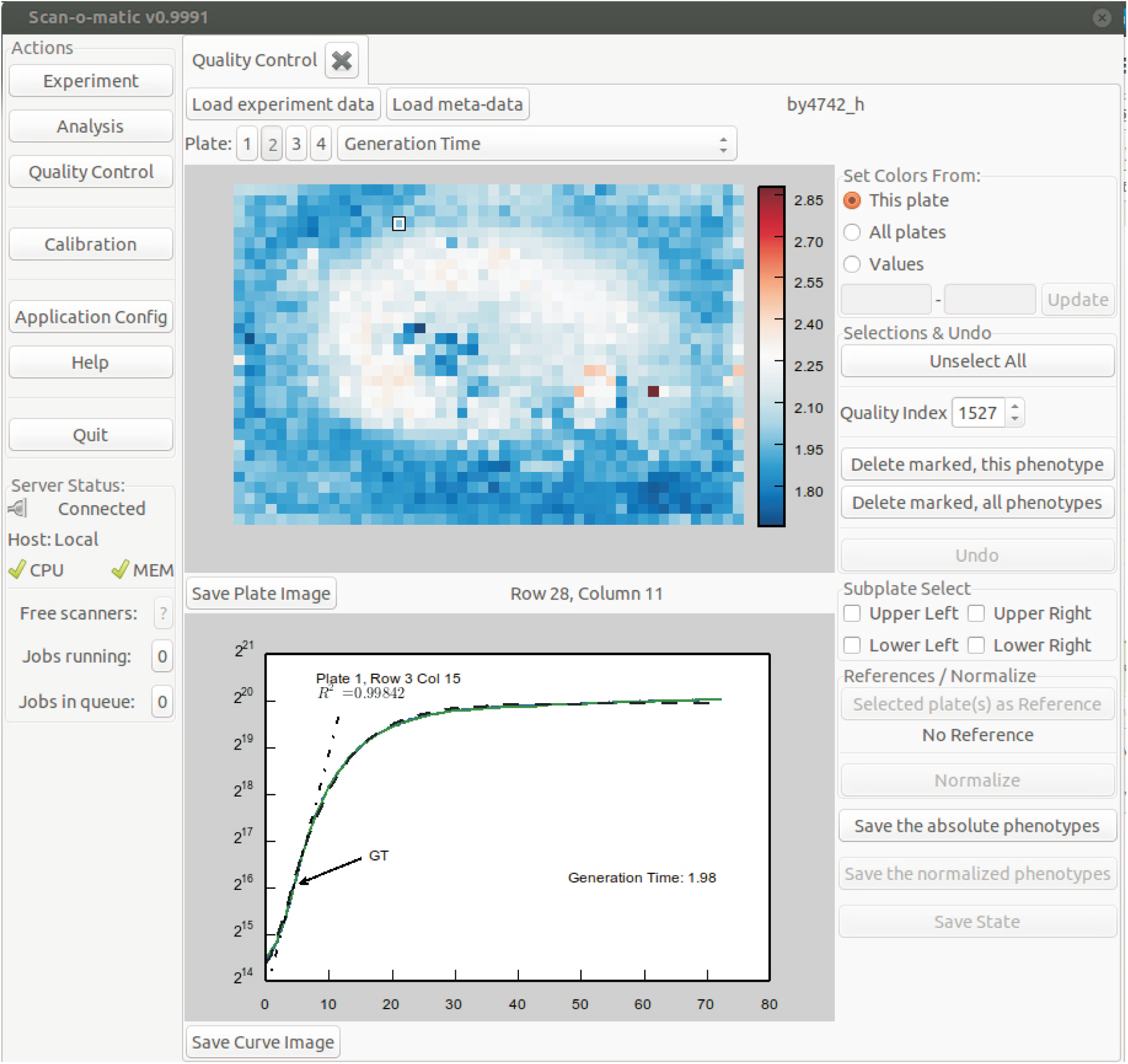
Post-experimental growth curve quality control interface. To give a visual representation of the distribution of phenotypes over a plate, we designed a user interface that shows each plate as a two-dimensional heatmap. The coordinates reflect colony positions and the colour intensity reflect the phenotype value. The phenotype to display is selected from a drop-down menu. In the lower part of the display, the currently selected colony’s growth curve is displayed together with phenotype information. The plate selector allows toggling between the four plates of the fixture, one at a time. The phenotype drop down selector allows selection of which phenotype to display in the heatmap format. The key output phenotype is the generation time/population size doubling time. Other phenotypes are: “Error of GT”, i.e. the error of the linear regression at the time of the population size doubling time extraction, the “Fit of the modified Chapman-Richards Model” and “time at maximal growth rate”. A large error of the linear regression, a poor model fit and a very early or very late extraction of population size doubling times suggest growth curves of low quality. Such suspected low quality growth curves can and should be manually inspected by selecting individual positions on the heatmap. Scan-o-matic will automatically display the colonies in ascending quality order to ease inspection. Low quality growth curves should be discarded before normalization, not to influence the normalization procedure. We urge users to be conservative in retaining growth curves, in particular if replication is high.

**Fig S9.**
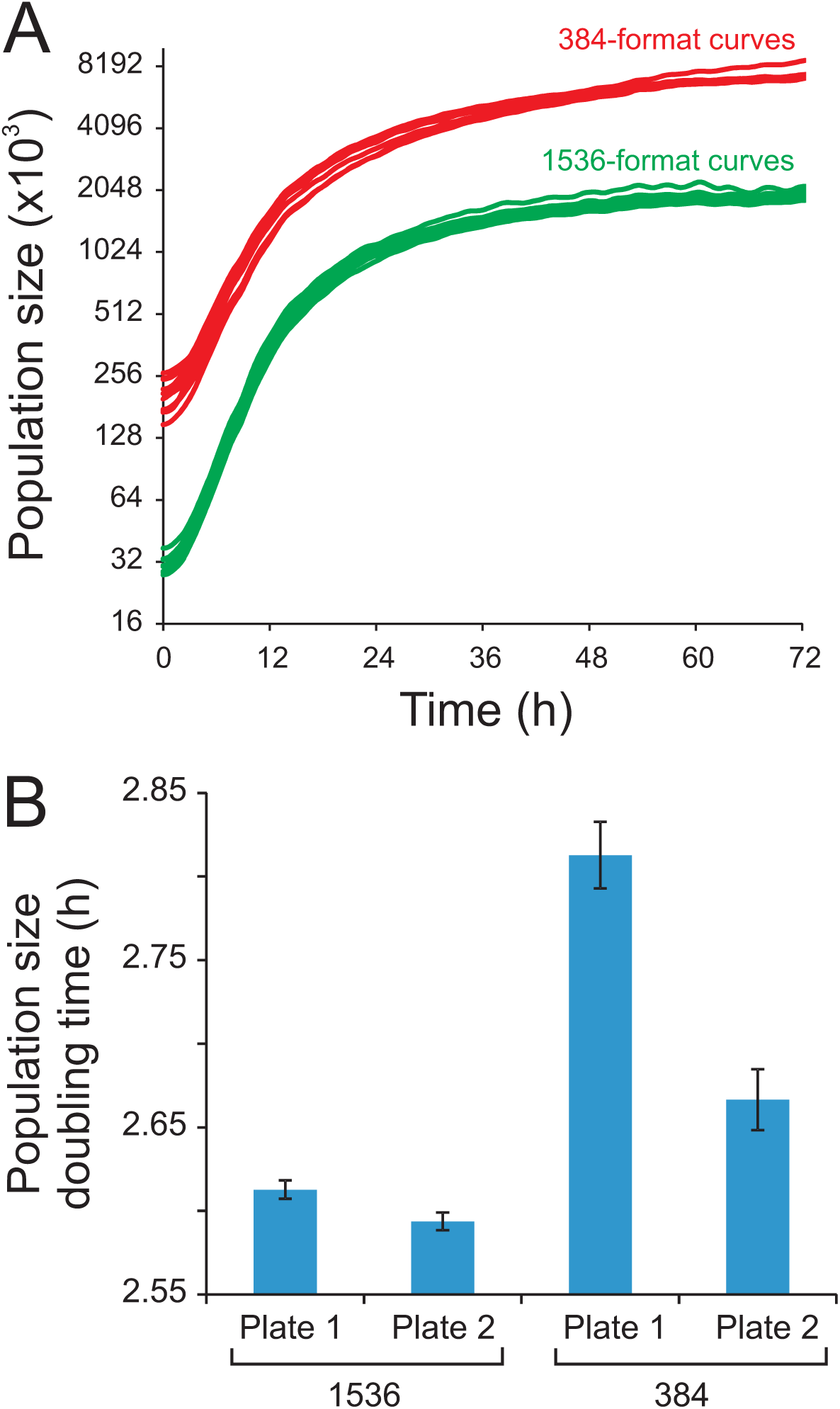
Effect of pinning format on Scan-o-matic growth curves. Genetically identical (WT, BY4741) colonies were pinned with either 1536 or 384 pins onto four identical no stress plates (synthetic defined media) with either 1536 (n=2 plates) or 384 pins (n=2 plates). Note that variation as a fraction of the signal, or CV, was much smaller for the 1536 (mean CV of 4.9% for 1536 vs. 8.2% for 384) A) Random growth curves from 1536 (n=10) and 384 pinned plates respectively (n=10). B) Mean population doubling time for each plate. Error bars = SEM (n = 1536 or 384).

**Fig S10.**
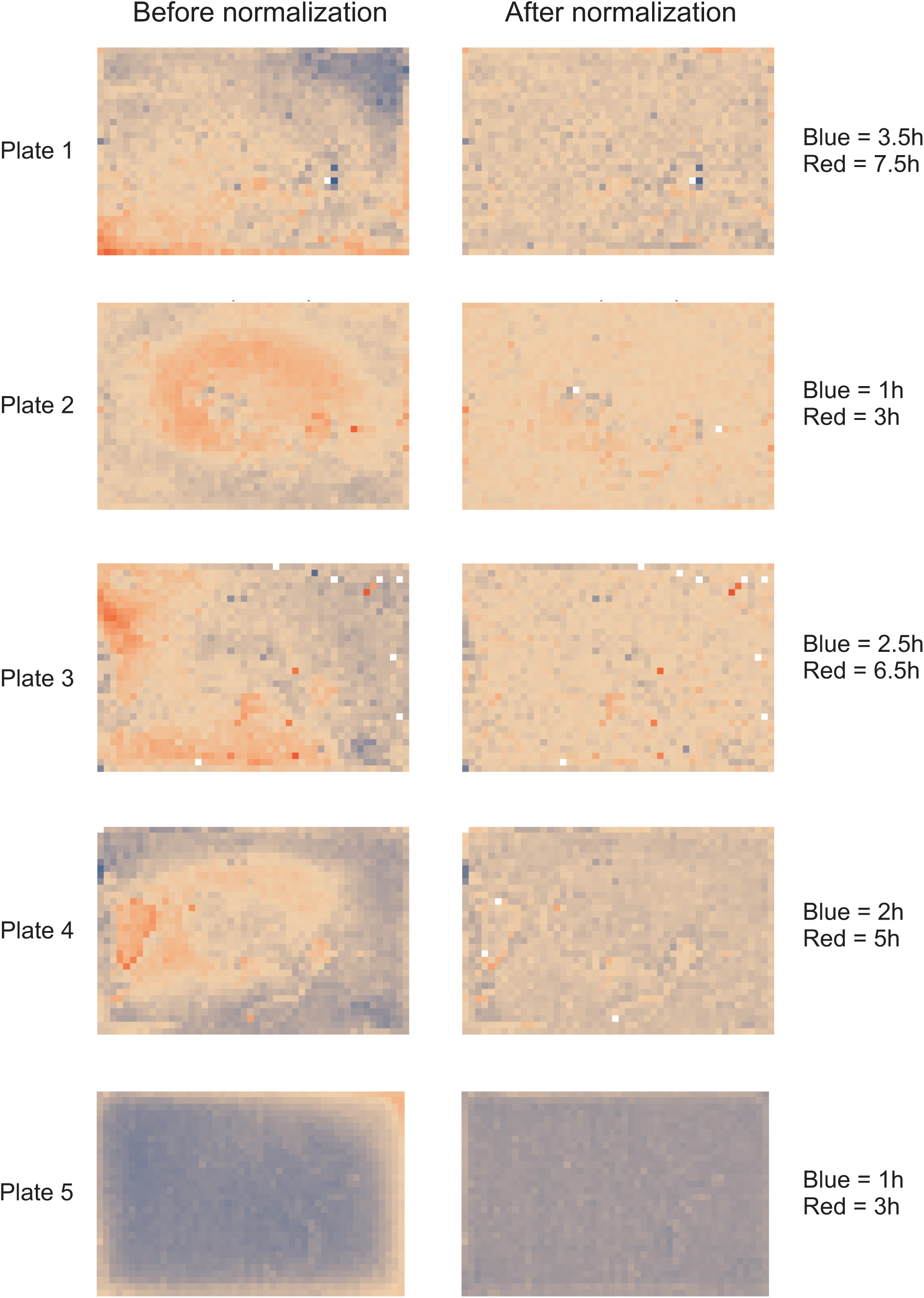
Spatial bias before and after reference grid normalization. Spatial bias is removed by reference grid normalization. Genetically identical reference colonies are pinned into every fourth colony position, creating a matrix of 384 control colonies on which a normalization surface of population doubling times is based. The local normalization surface is subtracted from each observation. Left panel = distribution of population size doubling times of 1536 genetically identical colonies across a plate, before normalization. Each square corresponds to a colony position. Right panel: As left panel, but color represents population size doubling times after reference grid normalization. Normalization was achieved by designating every 4^th^ position as a control position. Plates correspond to distinct environments, but all experiments are genetically identical, thus an even surface is expected. Color indicates population size doubling time with red = slow growth and blue = fast growth. Color scales are linear but ranges vary between plates to maximize resolution. Ranges are indicated to the right.

**Fig S11.**
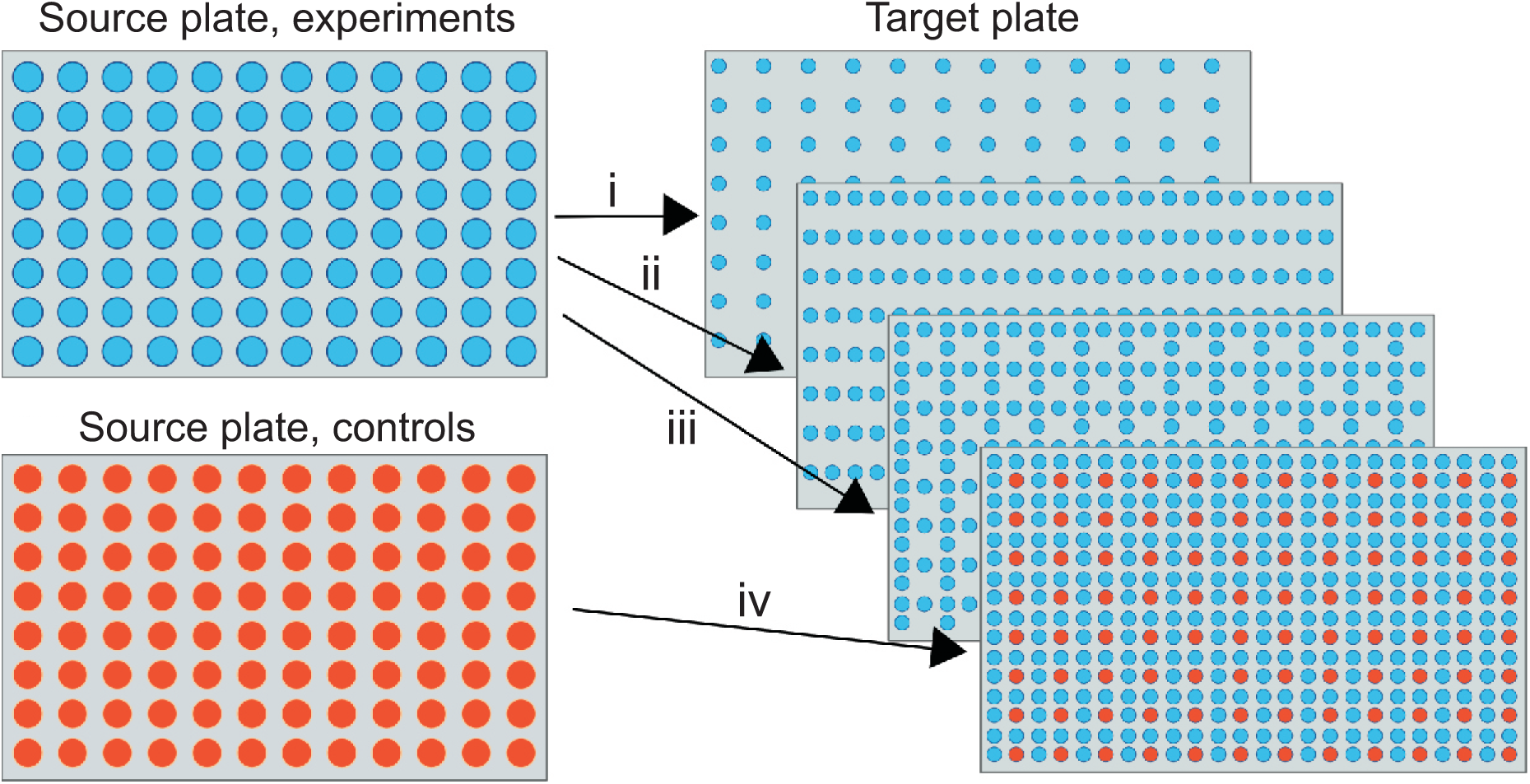
Introducing a reference grid of controls using a custom designed Scan-o-matic pinning program. Strains from two source plates, plate A containing intended experiments and plate B containing intended identical reference strains in all positions, are successively transferred to one target plate, C. Pinning is iterated x4 (i - iv) resulting in a 3:1 ratio of experiments relative controls on the target plate. Observe that plate C should not be used directly as experiment plate because the spatial bias of control colony growth on this plate poorly reflects the spatial bias of the growth of experiments. Instead, plate C should be used as a pre-culture for the real experimental plate.

**Fig S12.**
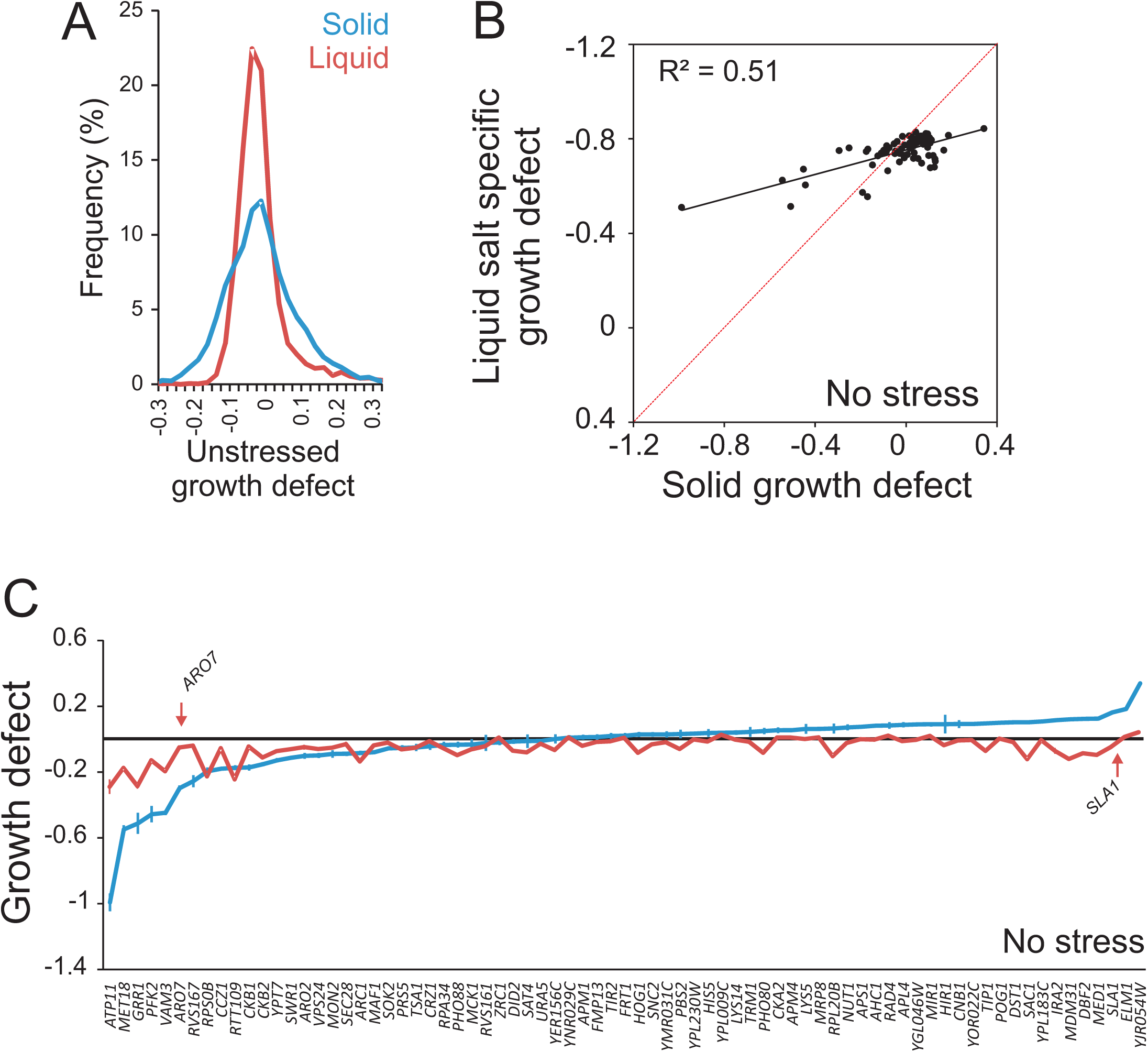
Scan-o-matic performance of gene deletion strains in absence of stress. The haploid *MAT***a** yeast deletion collection was cultivated in Scan-o-matic in absence of stress. Log_2_ population doubling times relative the control surface of WT controls were extracted. Negative values represent growth defects. A) Frequency distributions of salt-specific deletion strain growth effects, obtained by solid substrate cultivation in Scan-o-matic and by liquid microcultivation in a Bioscreen C. B-C) A subset of 70 deletion strains were re-cultivated in absence of stress at high replication, using Scan-o-matic (solid; n=24) and liquid (n=6) microcultivation respectively. Re-cultivations were performed in parallel, removing all conceivable systematic variation beside cultivation method. B) Growth effects of gene deletions in solid (Scan-o-matic) and liquid microcultivation. Regression (black, Pearson *R^2^* is indicated) and 1:1 lines (red) are shown. C) Gene deletion strains were ranked based on growth effects during solid substrate cultivation and growth effects were plotted. Error bars = SEM.

**Fig S13.**
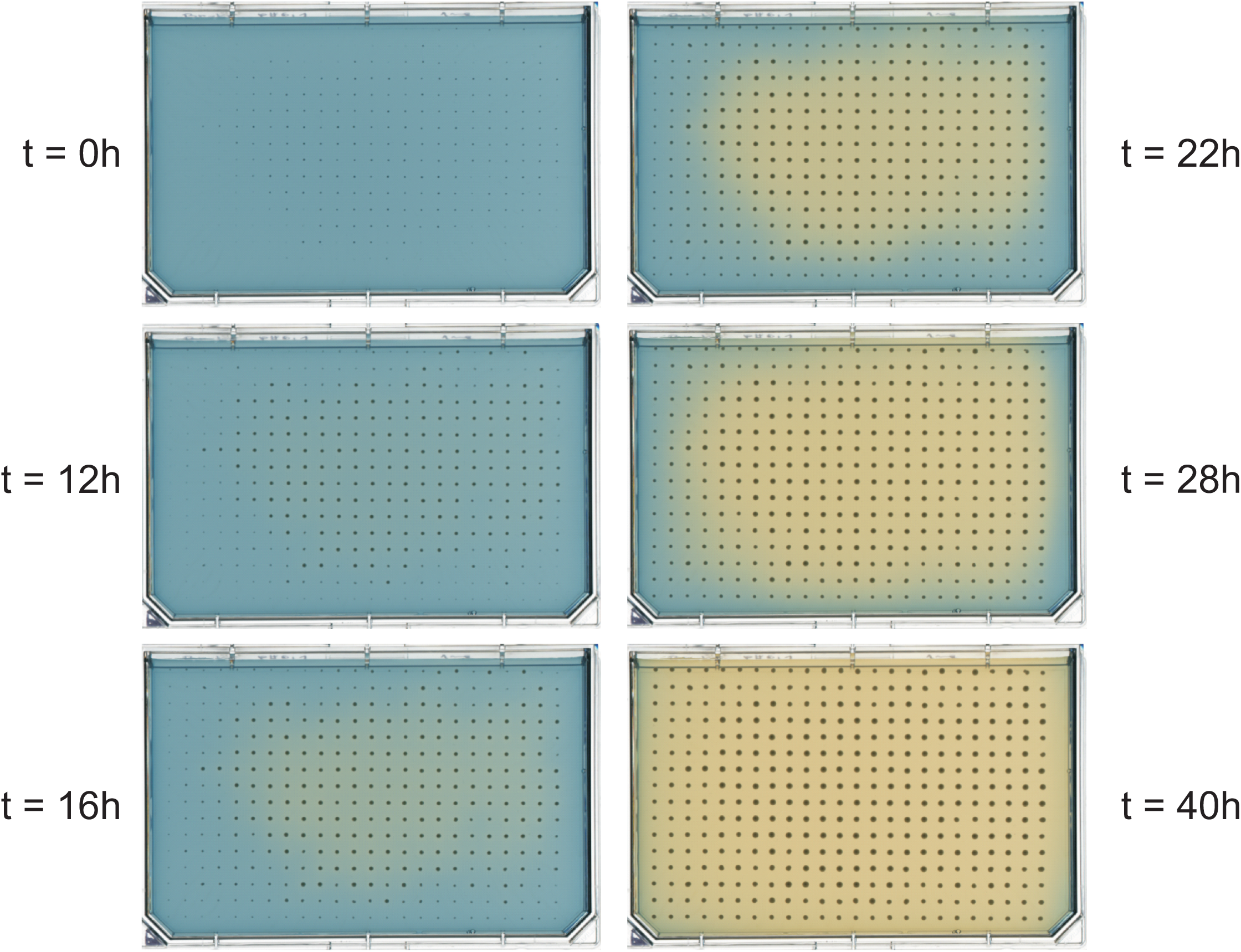
Spatial bias in the form of local changes in pH across solid media plates. Cells secrete organic acids metabolic by-products of carbon metabolism, lowering the pH of the local environment. Secreted acids diffuse slowly through the media, creating systematic spatial variations in pH as a function of colony size and colony metabolic state. External pH affects growth through altering a wide range of molecular phenotypes. Figure shows a time resolved view of pH change across a plate as a function of time. A plate was cast with unbuffered SC medium (initial pH = 6.0) supplemented with a pH indicator (1mg/50mL bromocresol green) and seeded with genetically identical BY4741 colonies (*his3Δ::kanMX4*) at uneven initial population sizes. Intense yellow = pH below 3.8, intense blue = pH above 5.4.

